# Modeling site-specific nucleotide biases affecting Himar1 transposon insertion frequencies in TnSeq datasets

**DOI:** 10.1101/2021.07.01.450749

**Authors:** Sanjeevani Choudhery, A. Jacob Brown, Chidiebere Akusobi, Eric J. Rubin, Christopher M. Sassetti, Thomas R. Ioerger

## Abstract

In bacterial TnSeq experiments, a library of transposons insertion mutants is generated, selected under various growth conditions, and sequenced to determine the profile of insertions at different sites in the genome, from which the fitness of mutant strains can be inferred. The widely used Himar1 transposon is known to be restricted to insertions at TA dinucleotides, but otherwise, few site-specific biases have been identified. As a result, most analytical approaches assume that insertion counts are expected a priori to be randomly distributed among TA sites in non-essential regions. However, recent analyses of independent Himar1 Tn libraries in *M. tuberculosis* have identified a local sequence pattern that is non-permissive for Himar1 insertion. This suggests there are site-specific biases that affect the frequency of insertions of the Himar1 transposon at different TA sites. In this paper, we use statistical and machine learning models to characterize patterns in the nucleotides surrounding TA sites associated with high and low insertion counts. We not only affirm that the previously discovered non-permissive pattern (CG)GnTAnC(CG) suppresses insertions, but conversely show that an A in the -3 position or T in the +3 position from the TA site encourages them. We demonstrate that these insertion preferences exist in Himar1 TnSeq datasets other than *M. tuberculosis*, including mycobacterial and non-mycobacterial species. We build predictive models of Himar1 insertion preferences as a function of surrounding nucleotides. The final predictive model explains about half of the variance in insertion counts, presuming the rest comes from stochastic variability between libraries or due to sampling differences during sequencing. Based on this model, we present a new method, called the TTN-Fitness method, to improve the identification of conditionally essential genes or genetic interactions, i.e., to better distinguish true biological fitness effects by comparing the observed counts to expected counts using a site-specific model of insertion preferences. Compared to previous methods like Hidden Markov Models, the TTN-Fitness method can make finer distinctions among genes whose disruption causes a fitness defect (or advantage), separating them out from the large pool of non-essentials, and is able to classify the essentiality of many smaller genes (with few TA sites) that were previously characterized as uncertain.

## Introduction

TnSeq has become a popular tool for evaluating gene essentiality in bacteria under various conditions (Cain, Barquist et al. 2020). The most widely used transposons for bacterial TnSeq are those in the *mariner* family, such as Himar1 (Sassetti, Boyd et al. 2003). To date, it has generally been assumed that the Himar1 transposon, frequently used to generate the transposon libraries, inserts randomly at TA dinucleotide sites in non-essential regions across the genome (Lampe, Akerley et al. 1999). The abundance of transposon insertions at each TA site can be quantified efficiently using next-generation sequencing (Long, DeJesus et al. 2015). Genes or loci with an absence of insertions are considered to be essential, as disruption in these regions are not tolerated (Sassetti, Boyd et al. 2003). Genes or loci with a reduced mean insertion count are considered mutants with growth defects, as disruptions in these regions are not fatal but cause growth impairments or fitness defects (van Opijnen, Bodi et al. 2009). Genes that have significant changes in mean counts between conditions are deemed as conditionally essential (Gawronski, Wong et al. 2009).

There are several sources of noise in TnSeq experiments, including stochastic variations in the library generation process as well as instrument and sampling-error in DNA sequencing, resulting in a high variability in insertion counts. Statistical methods developed thus far to assess gene essentiality typically assume that insertions occur randomly at TA sites in non-essential regions, and the reason some sites have more insertions than others is largely due to stochastic differences in abundance in the library. However, some studies suggest that transposon insertions at non-essential sites is influenced by the surrounding nucleotides or genomic context. Transposons Tn5 and Mu (not restricted to TA dinucleotides) showed a bias towards insertions in GC-rich regions and resulted in a less uniform distribution of insertions in the A-T rich genome (61% AT) of *C. glabrata* than their notably less-biased counterpart Tn7 (Green, Bouchier et al. 2012). In addition, Lampe (Lampe, Akerley et al. 1999) showed that local bendability of the DNA strand can affect the probability of Himar1 insertion at different chromosomal locations in *E. coli*. Furthermore, an analysis of 14 independent transposon libraries in *M. tuberculosis* (Mtb) H37Rv identified a local sequence pattern around certain TA sites that was non-permissive for Himar1 insertions (CG)GnTAnC(CG) (DeJesus, Gerrick et al. 2017). This sequence pattern extended to ∼9% of sites in non-essential regions which almost always had counts of zero (DeJesus, Gerrick et al. 2017).

In this paper, we use statistical and machine learning models to identify patterns in the nucleotides surrounding TA sites associated with high and low insertion counts. We discover nucleotide biases within a ±4-base pair window around the TA site that suppress Himar1 insertions, and other patterns that appear to select for them (i.e., associated with high insertion counts). We capture these biases in a predictive model of Himar1 insertion preferences that can be used to predict expected insertion counts at any TA site as a function of the surrounding nucleotide context. We demonstrate that these insertion preferences exist in other Himar1 TnSeq datasets from *Mtb*, as well as other mycobacterial and non-mycobacterial species. The final predictive model explains about half of the variance in insertion counts, presuming the rest comes from stochastic variability between libraries or due to sampling differences during sequencing. We demonstrate that this model can be used to improve the assessment of genes’ fitness by comparing the observed counts to expected counts using a site-specific model of insertion preferences.

## Results

### Insertion counts at TA sites are correlated between libraries

Variability in insertion counts at TA sites can be attributed to various sources, including abundance in library, experimental randomness, and local sequence biases, as well as genuine biological significance (fitness effects). To attempt to differentiate these, we re-analyzed a previously published collection of 14 independent Himar1 TnSeq libraries grown in standard laboratory medium (DeJesus, Gerrick et al. 2017). An extended HMM analysis the 14 datasets (see Methods) suggests that approximately 11.6% of the organism’s TA sites are essential for growth, and insertions in approximately 3.5% of the sites can cause a growth defect. In addition, 9% of sites in non-essential regions have few to no insertions due to a non-permissive sequence pattern (DeJesus, Gerrick et al. 2017). Insertions at TA sites in regions other than these are generally expected to occur randomly. If true, the insertion counts at the same TA site in different libraries would be expected to be uncorrelated on average. However, our analysis of the 14 H37Rv Tn libraries shows that there is substantial correlation of counts at individual TA sites, suggesting that some TA sites have a higher propensity for Himar1 insertion than others. Figure 1 shows the distribution of log_10_ insertions counts from each library in a genomic region with 75 TA sites (the log of counts was taken to better fit a Gaussian distribution). Each library was TTR-normalized to make counts from datasets of different total size comparable (DeJesus, Ambadipudi et al. 2015). Panel A shows that mean insertion counts differ widely among non-essential TA sites, and the variability between TA sites is more than within each site. Thus, high counts at a TA site in one library tend to have high counts in other libraries and similarly, sites with low counts occur symmetrically across the libraries. As a comparison, insertion counts at TA sites (excluding those marked essential or following the non-permissive pattern) were randomized within each library. Panel B shows the same 75 consecutive TA sites in this randomized dataset. When randomized, the distribution of counts at sites in non-essential regions is much more uniform. The average variance of all insertions within a TA site is 0.430, significantly lower (p-value < 0.001) than the variance of 0.929 found in the randomized dataset. This makes it evident that the correlation of log insertion counts across libraries is greater than expected. In fact, pairwise correlations of the randomized datasets range from 0.15 to 0.33, averaging to 0.28. Pairwise correlations of 14 libraries are considerably higher, ranging from 0.5 to 0.97, averaging to 0.62 (see Supplemental Figure S1). 80 of the 90 pairwise correlations had significant p-values (< 0.01) from comparison by a two-tailed t-test (see Supplemental Table T1). A significant high correlation across libraries suggests there are site-specific influences, in addition to those previously observed, on insertion probabilities at different TA sites.

**Figure 1:**
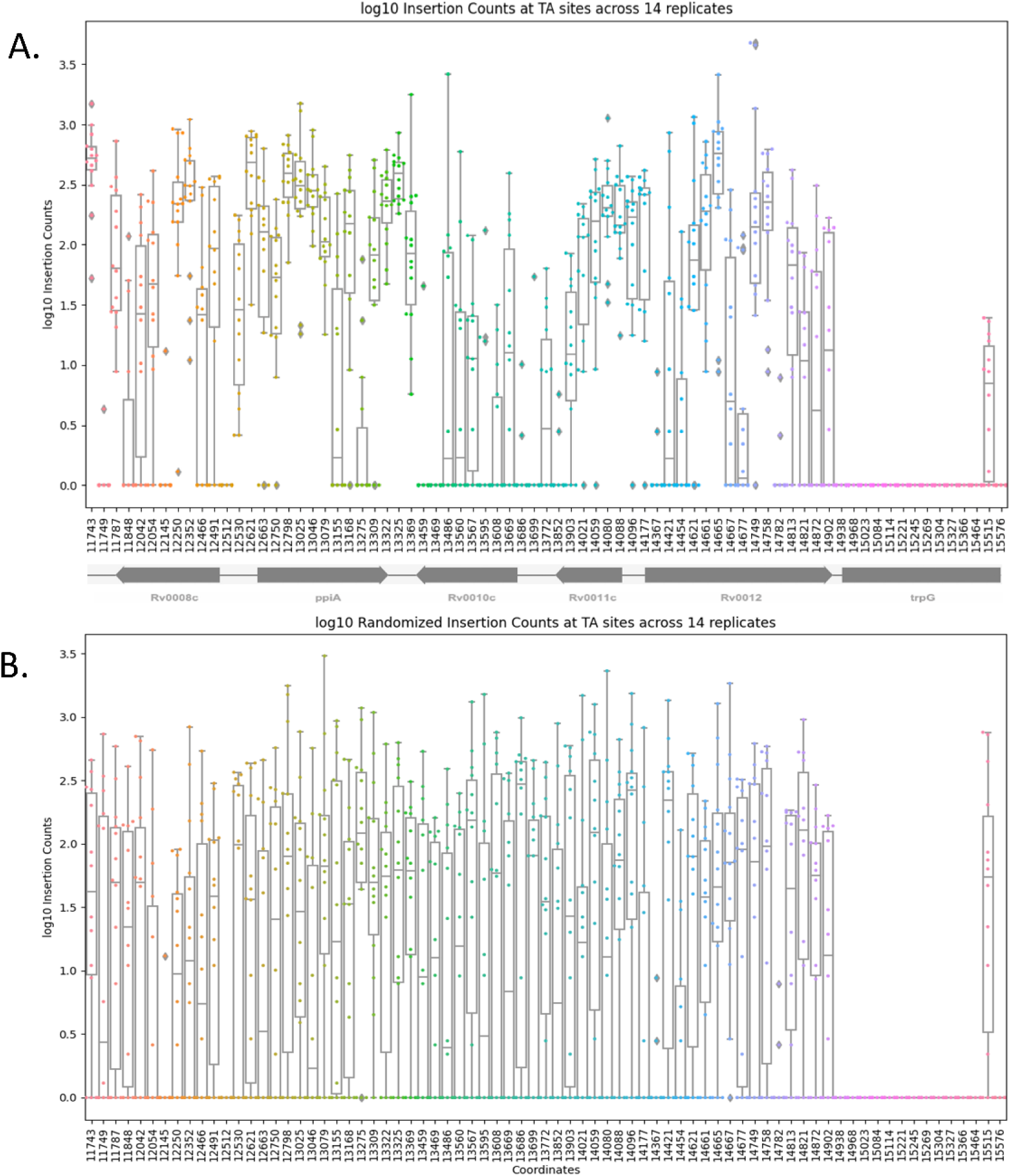
Insertion Counts across 14 H37Rv libraries in a region spanning 75 consecutive TA sites. In **Panel A**, a point is plotted for insertion counts at each coordinate for each replicate. This scatter plot is then overlayed with a box-and-whisker plot reflecting the mean and range of insertion counts at each site. The region includes *trpG* for comparison, which is an essential gene, and hence insertion counts are 0 in this gene. In the non-essential genes, the insertion counts vary more between TA sites than within, supporting that some TA sites have a higher propensity for insertions than others. **Panel B** shows the same 75 sites after randomizing the insertion counts at all TA sites except those marked ES and those showing the non-permissive pattern. The mean and range of counts at each non-essential TA site are much more uniform when randomized.

### Modeling Insertion Counts using Linear Regression

To determine whether the nucleotides surrounding a TA site influence the probability of insertion, we examined the association of proximal nucleotides on insertion counts, averaged over all non-essential TA sites in the genome. Figure 2 presents evidence of site-specific nucleotide effects that influence the relative abundance of insertions at TA sites. Panel A shows overall nucleotide probabilities ±20 bp from the TA site. Most of the deviation in nucleotide probabilities occurs within 4 bp of the central TA site, with probabilities varying up to 20% for some nucleotides. Further insight can be gained by dividing the TA sites into thirds: sites with lowest counts, sites with medium counts, and sites with highest counts. Panel B, depicting the lowest third of the range of insertion counts, shows an increase in probabilities of nucleotides C and G and a decrease in probabilities of nucleotides ‘A’ and ‘T’ especially at positions ±2 and ±3. Panel D, depicting the highest third of insertion counts, also shows drastic changes in nucleotide probability, with a notable increase in propensity for ‘A’ at -3 and ‘T’ at +3. These observations suggest a correlation between the magnitude of insertion counts and nucleotides surrounding TA sites. Thus, insertion counts at a TA site could be affected by the surrounding nucleotides.

**Figure 2:**
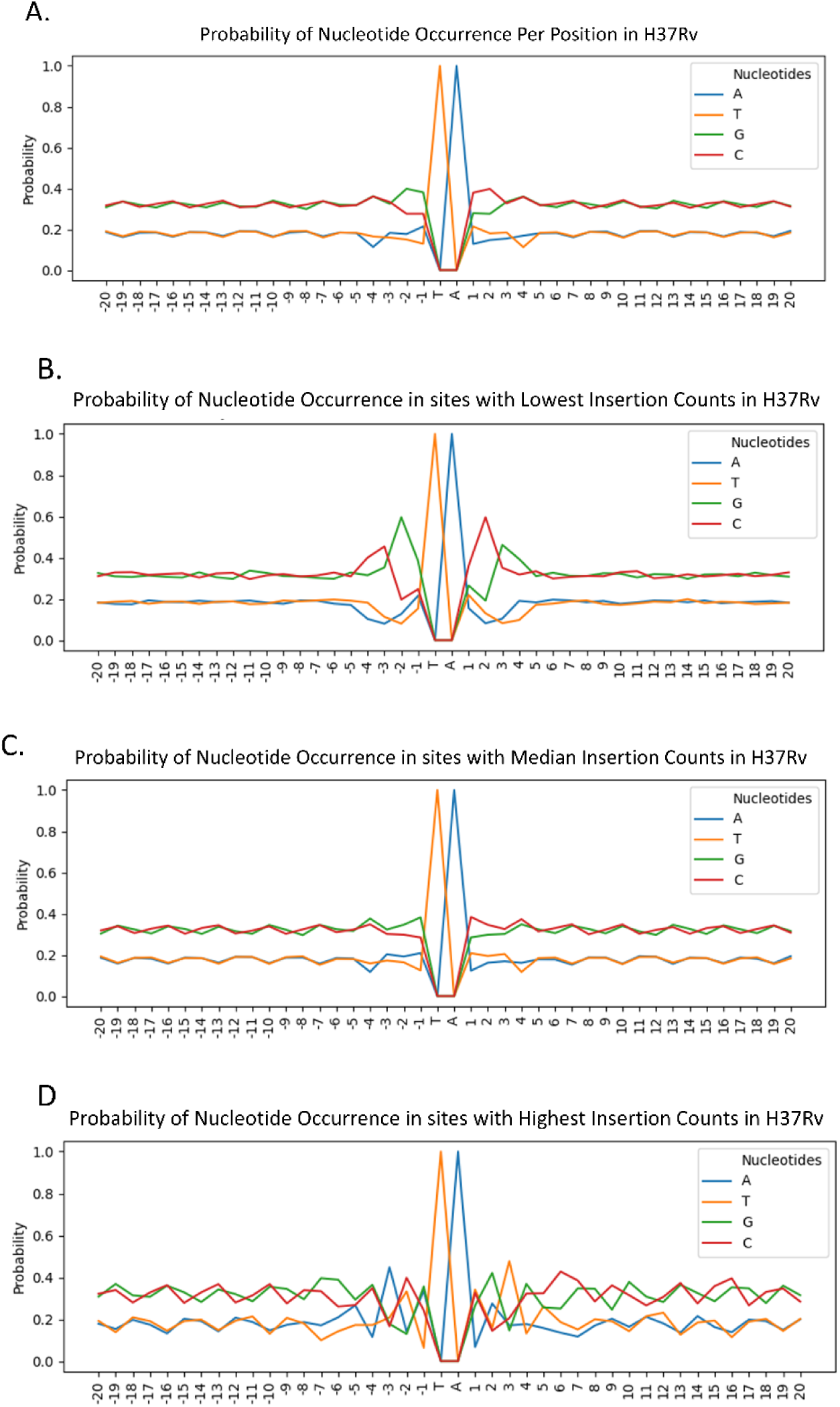
Nucleotide Probabilities at positions -20…+20 from the TA site, for three ranges of Insertion Counts. The 3 ranges of the log insertion counts depicted in Panels B,C,D were found by dividing the difference in the maximum log count (10.83) and minimum log count (-2.30) by 3. The boundaries of the splits were at 6.45 and 2.07. **Panel A** shows the pattern across all ∼65000 TA sites in non-essential regions. **Panel B** shows the pattern across 8992 sites in the lower third of the range, **Panel C** shows the pattern across 51164 sites in the middle third of the range and **Panel D** shows the pattern across 1172 sites in the higher third of the range.

We trained a linear regression model on the 40 nucleotides surrounding the TA site (positions - 20…+20) to predict insertion counts in known non-essential regions (67,670 TA sites) using the mean counts from the 14 libraries of H37Rv. The input to the model was a one-hot-encoding of the nucleotides, where each nucleotide at each position was represented by 4 bits and concatenated into a bit vector, totaling 160 binary features. The resulting linear model was:

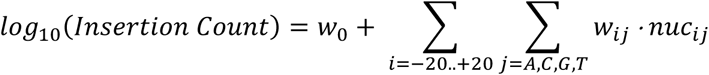

where nuc_ij_=1 if nuc[i] = j, nuc[i] is the nucleotide at position i relative to the TA site and weights w_ij_ correspond to each of the 160 binary features. This formula is equivalent to a dot-product a 160-bit vector (plus an intercept) with a vector of weights, log_10_ (Insertion Count) = w_0_ + w_1_ b_-20=A_ + w_2_ b_-20=C_ +w_3_ b_-20=T_ + w_4_ b_-20=G_ +…+ w_157_ b_+20=A_ + w_158_ b_+20=C_ +w_159_ b_+20=T_ + w_160_ b_+20=G_, where every four bits encode the nucleotide at a position ± 20 bp from the TA site. The model was trained and evaluated using 10-fold cross validation. Figure 3 shows the average correlation between predicted and observed log_10_ insertion counts. The model has some predictive power (R^2^ value of 0.32), but also has high variance. A slight bias can be seen in the figure, where the low counts are predicted too high, and the high counts too low. This is a consequence of the regression model making predictions that do not span as wide of a range as the actual data, due to inaccurate predictions for the sites with the most extreme (largest or smallest counts). The accuracy of predictions made by this initial simplified regression will increase with improved models (below) and thus this effect will be reduced.

**Figure 3:**
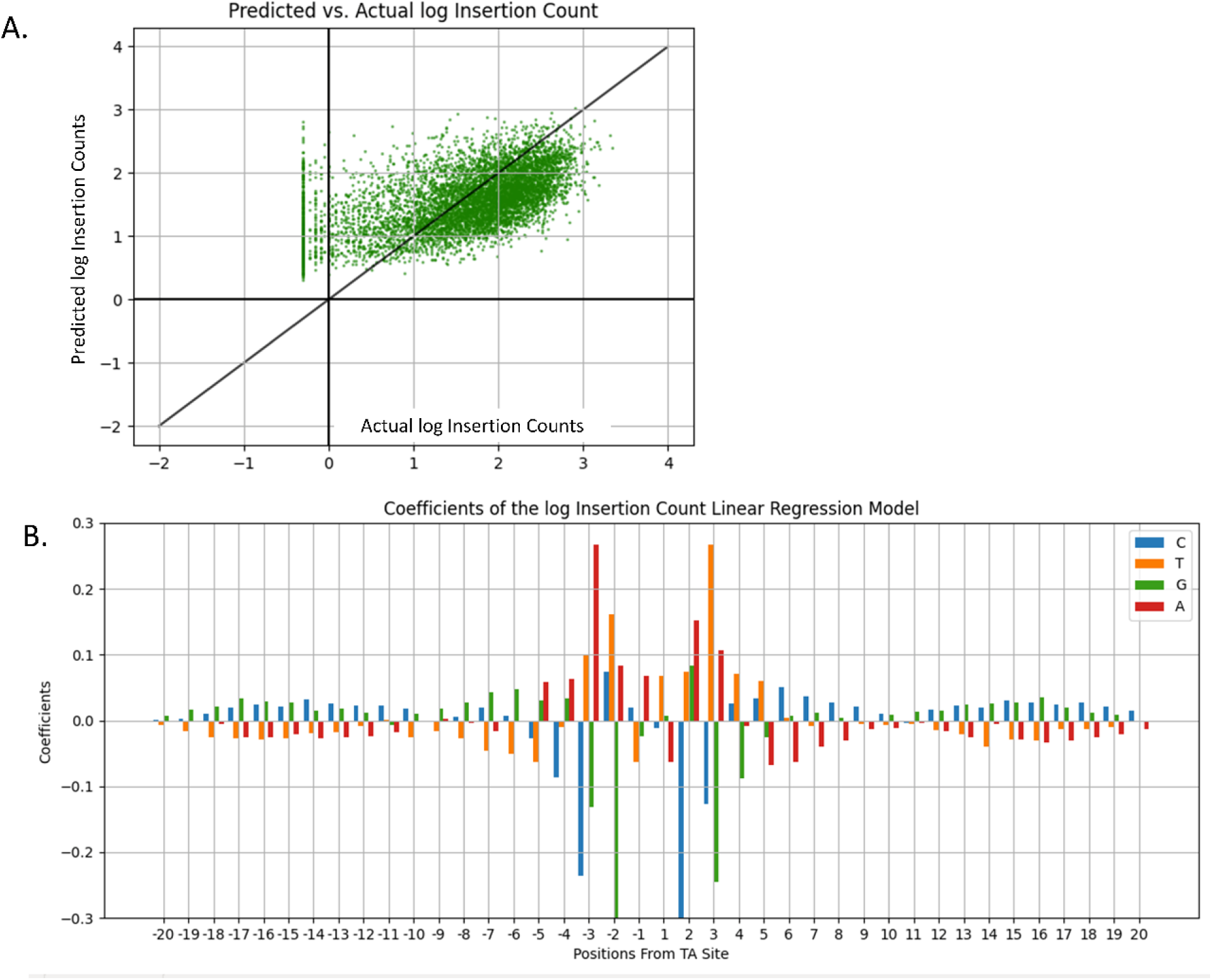
Coefficients and Accuracy Assessment of Linear Regression Model trained on nucleotides as covariates. **Panel A** shows Predicted Counts vs. Actual log Insertion Counts using Linear Regression. The average predictive power of the Linear Regression Model trained with one-hot-encoded nucleotides as the input and log insertion counts as the output using 10-fold cross validation. The predictive power is moderate (R^2^=0.318), meaning it is able to explain 31% of the variation in insertion counts based on surrounding nucleotides. **Panel B** shows Coefficients from the trained Linear Model. The coordinates along the x-axis give the positions relative to, but not including, the TA site. The model is trained on one-hot-encoded nucleotides and a target value of log Insertion Counts. The symmetry of the pattern is visible in positions -4, -3, -2, -1 and +1, +2, +3, +4. The non-permissive pattern (CG)GnTAnC(CG) is visible in this window, as well as high coefficients associated with “A” and “T”.

In Figure 3B, nucleotides with highest coefficients in the trained model are located within a window of ±4 bp around the TA site. The pattern created by the nucleotides of these coefficients are consistent with the non-permissive pattern (CG)GnTAnC(CG) previously reported (DeJesus, Gerrick et al. 2017). Nucleotide ‘G’ has the highest absolute coefficient value in the -2 position and ‘C’ has the highest absolute coefficient value in the +2 position. Moreover, both ‘C’ and ‘G’ have similarly high absolute coefficients in the -3 and +3 positions. In addition to the confirmation of the non-permissive pattern (large negative coefficients for ‘G’ at -3 and ‘C’ at +3), the figure shows nucleotides ‘A’ and ‘T’ with relatively high positive coefficients in positions -3 and +3 from the TA site. These patterns reinforce the observations made in Figure 1 and provide further evidence of previously undetected site-specific nucleotide biases that affect Himar1 insertion counts.

### Prediction of insertion counts at TA sites relative to local average counts

We assume that insertion counts are proportional to the permissiveness of a site i.e., a site with a less permissive pattern will have lower counts than a site with a more permissive pattern. However, insertion counts are also affected by biological fitness. It is likely that a TA site with a specific nucleotide pattern in a fitness-defect gene will have a lower insertion count than a TA site with the same pattern in a non-essential gene. But this effect (decrease or increase in counts) should be shared by multiple TA sites locally. We can compare the insertion count observed at a site to the observed counts at other TA sites in the region, the level of which should reflect the general fitness effect of disrupting the gene. Thus, modeling this relative (or local) change in insertion counts would allow us to factor out biological effects on counts and focus on the effect of nucleotide patterns on the insertion counts.

This change in insertion counts is quantified for every TA site as a log-fold-change (LFC) value. The local average was calculated for each site by taking the mean insertion counts from the previous 5 and next 5 TA sites from the site of interest (i.e., using a sliding window of 11 consecutive TA sites).

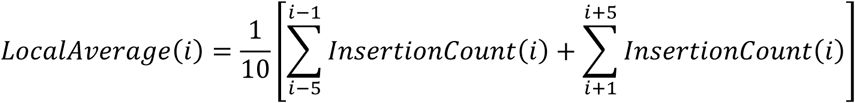

The local mean excludes the central site itself and any locations marked as essential during pre-processing. The LFC for each TA site was calculated by taking the log insertion count at that site plus a pseudo count of 10 (to smooth out high variability of LFCs for sites with low counts) and dividing it by the local average:

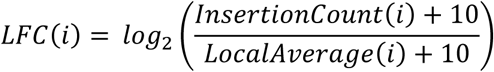

As with the previous model, this linear model was trained and tested using 10-fold cross validation. The resulting model (see Supplemental Figure S2) has an average R^2^ value of 0.38, indicating that training the model to predict changes in insertion counts (relative to local mean) rather than absolute insertion counts greatly reduces the noise due to local fitness effects (e.g., in a gene where insertion cause growth defects, systematically reducing abundance of insertions in the region). This allows the model to better capture the effect of nucleotides surround TA sites on Himar1 insertion preferences.

### A neural network model explains up to 50% of the variability in insertion counts

As they can capture non-linear patterns, neural networks are considered to be some of the most powerful predictors in Machine Learning (Rumelhart, Hinton et al. 1986). To see if we could increase the accuracy of our model, we tried using our data to train a fully connected multi-layer feed-forward Neural Network. The model contained one hidden layer of 50 nodes. This parameter along with other hyper parameters of the network were tuned using a grid search (see details in Methods and Materials). A random subset of 70% of the data was used to find the ideal hyper parameters through cross validation, with the remaining 30% of the data used to test the final hyper parameters. A 10-fold cross-validation of the entire dataset was used to train and test the model i.e., judge the model accuracy using our data. The input to the model consisted of bit-vectors encoding nucleotides surrounding each TA site in the dataset, totaling to 160 features. The target value was LFCs (log-fold-changes of insertion counts relative to local mean). The model performed better than the previous models with an average R^2^ of 0.51 (R^2^ of 0.509 on the hyper parameter test data) (see Supplemental Figure S3). Thus, the neural network can explain around half of the variability in insertion counts at TA sites based on surrounding nucleotides; presumably, the remaining differences in counts still reflect stochastic differences in abundance between libraries (or other influences on TA insertion preferences for which we have not yet accounted). However, as is typical for neural networks, this model (as a matrix of connection weights) does not provide us much insight into nucleotide patterns that led to the predictions for the TA sites.

### Certain Nucleotides Surrounding TA Sites are Associated with High or Low Insertion Frequencies

It has been previously noted that there are biases in distributions of nucleotides surrounding TA sites, making them more permissive or less permissive. If a site has a pattern that is considered more permissive, it should have a higher insertion count than its neighbors and thus a positive LFC. The opposite is true for sites with a less permissive pattern. They should have lower counts than their neighbors and thus negative LFCs. The heatmap in Figure 4 was generated to visualize any additional nucleotide biases that may result in unusually high or low insertion counts. For each nucleotide N and position P within ±20 bp of a TA site, the mean LFC was calculated over the subset of TA sites having nucleotide N at position P (Methods and Materials). The heatmap reinforces the idea illustrated in Figure 2 of the correlation between nucleotide biases and insertion count magnitudes. A ‘G’ in position -2 and its symmetric counterpart ‘C’ in position +2, as well as ‘C’ in the -3 position and its counterpart ‘G’ in +3 position are associated with low mean LFCs. This indicates that TA sites with at least one of these nucleotides in their relative positions tend to have lower insertion counts than their neighbors, consistent with the nucleotide bias represented by the non-permissive pattern (CG)GnTAnC(CG) observed in (DeJesus, Gerrick et al. 2017). Similarly, there is a distinctive pattern for positive mean LFCs: an ‘A’ in position -3 and its counterpart ‘T’ in position +3 are both associated with higher mean LFCs, and hence can be interpreted as being more permissive for Himar1 insertions (associated with increased counts). However, the effects of multiple biases appearing in a single sequence are not additive. For instance, a ‘C’ in the -2 and an ‘A’ in the -3 position do not “cancel” each other out; they are interdependent. We quantify how effects like these combine in the tetra-nucleotide model below.

**Figure 4:**
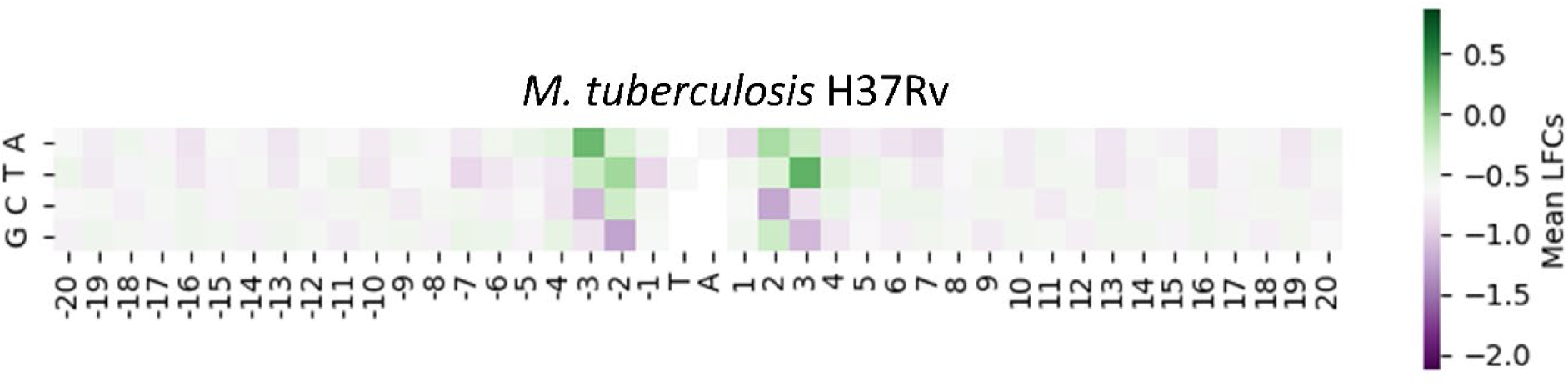
Enrichment and Depletion of Nucleotides surrounding TA sites for the 14 libraries of H37Rv in-vitro. The mean of each filtered nucleotide at every position in a 20-bp window of the TA site in H37Rv dataset is visualized here. The heatmap centered at the median of mean LFCs calculated with a median +1.5 max and median-1.5 minimum. Nucleotides colored green at certain positions are enriched, while those colored purple are depleted (relative to the global average nucleotide content).

There appears to be a slight periodic pattern of the G and C nucleotides surrounding the TA site, between 20 and 4 bp from the TA site in Figure 4 (also evident in Figure 2). The nucleotides show an increase in mean LFC for every third position in the sequence. Representing this pattern in a simplistic manner and comparing it to the LFC target variable showed little correlation. Thus, this periodic sequence was not incorporated in our model.

### Symmetric Tetra-nucleotide Linear Model (STLM)

To gain more insight into the nucleotide patterns observed through the heatmaps, we devised a variant of the linear model, called the STLM (Symmetric Tetra-Nucleotide Linear Model). In the linear models previously mentioned, the pattern associated with individual nucleotide was implicitly assumed to be additive, and thus each nucleotide position was treated as an independent variable. But we wondered whether a stronger pattern may be found through combinations of these nucleotide positions, which can represent non-linear interactions.

Training the linear model to predict LFCs based only on the nucleotides in a window of ±4 bp from the TA site yielded nearly identical results to the regression predicting LFCs using all 40 nucleotides (R^2^ = 0.35 and the same coefficient pattern for nucleotides in range -4…+4) indicating that most of the influence on LFC predictions is within an 8 bp window (see Supplemental Figure S4). This is reinforced by the heatmaps, as a majority of the apparent effects occur within 4 bp from the TA site. If we use all the sequence combinations of the nucleotides in positions -4…+4 as features in our model, we will have 4^8^ =65,536 features (i.e., terms in a linear model, or inputs to a neural network). However, the patterns of nucleotide biases are symmetrical (reverse-complement), as shown by the heatmaps, thus making the distinction between all 8 nucleotides unnecessary. The four nucleotides upstream of the TA appear to affect the insertion counts in the same way as the reverse-complement of the 4 nucleotides downstream of the TA site. Therefore, it is only necessary to capture the association of 4 nucleotides at a time on LFCs in the model. Hence, we shift to training our models based on combinations of 4 nucleotides, i.e., tetra-nucleotides, which reduces the number of features in our model to 4^4^ = 256.

As input to the STLM, each TA site is represented as a vector where all features are set to 0 except for the upstream tetra-nucleotide and reverse-complemented downstream tetra-nucleotide (see Figure 6). This is essentially the same as adding two bit-vectors, one vector with the bit for the upstream tetra-nucleotide on and another separate vector with the bit for the downstream tetra-nucleotide on. The result is a sparse 256-bit vector with only 2 bits on (except when the two tetra-nucleotides are the same, in which case the single feature value for the tetra-nucleotide is set to 2). The result is a linear model that follows the equation

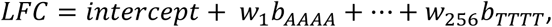

where *w*_1_… . *w*_256_are the weights associated with tetra-nucleotides (to be trained by the model) and *b_AAAA_*… *b_TTTT_* are the bits corresponding to the presence of the adjacent tetra-nucleotide features for every TA site. Encoding both the upstream and reverse-complemented downstream tetra-nucleotides allows us to use the same model to represent the bias from both sides of the TA simultaneously as independent features, additively contributing equal weight. Assume for a given TA site, both upstream and downstream tetra-nucleotides are associated with high LFCs; then they will reinforce to predict an even higher insertion count for that site. But if the upstream tetra-nucleotide has a trend to contribute a high LFC and the reverse-complemented downstream tetra-nucleotide has a trend to contribute a low LFC, they will tend to cancel each other out.

As seen in Figure 5A, 10-fold cross validation using the H37Rv data resulted in an average R^2^ value of 0.469. This R^2^ value is slightly lower than, but nearly equal to, that of the neural network (p-value < 0.01 from two-tailed t-test). However, the STLM provides us more insight into patterns contributing to the prediction of the LFCs. In a regression with these tetra-nucleotide features, we expect each coefficient (i.e., weights) of the model to correlate with the average LFC associated with each tetra-nucleotide (over TA sites surrounded by these tetra-nucleotides). Figure 5B shows the relationship of the STLM coefficients, and the mean observed LFCs of the corresponding tetra-nucleotides (shifted on the y-axis by the bias (intercept) in our data). The strong linear trend visible adheres to the expectation of a high correlation and indicates our model accurately represents our data. The individual tetra-nucleotide coefficients are shown in Panel C, sorted in decreasing order (See Supplemental Table T4 for full table). Consistent with the patterns observed in the heatmaps in Figure 4, the bottom ten features associated with low coefficients (predictive of low mean LFCs) all have a ‘G’ in the 2nd position upstream of the TA sites and a ‘C’ or ‘G’ in the 3rd position. The features associated with the top ten coefficients, thus higher LFC values, all have an ‘A’ in 3rd position upstream from the TA site. However, the strength of the STLM is that it accounts for combinations of 4 nucleotides together at a time, resolving cases where single-nucleotide patterns might conflict.

**Figure 5:**
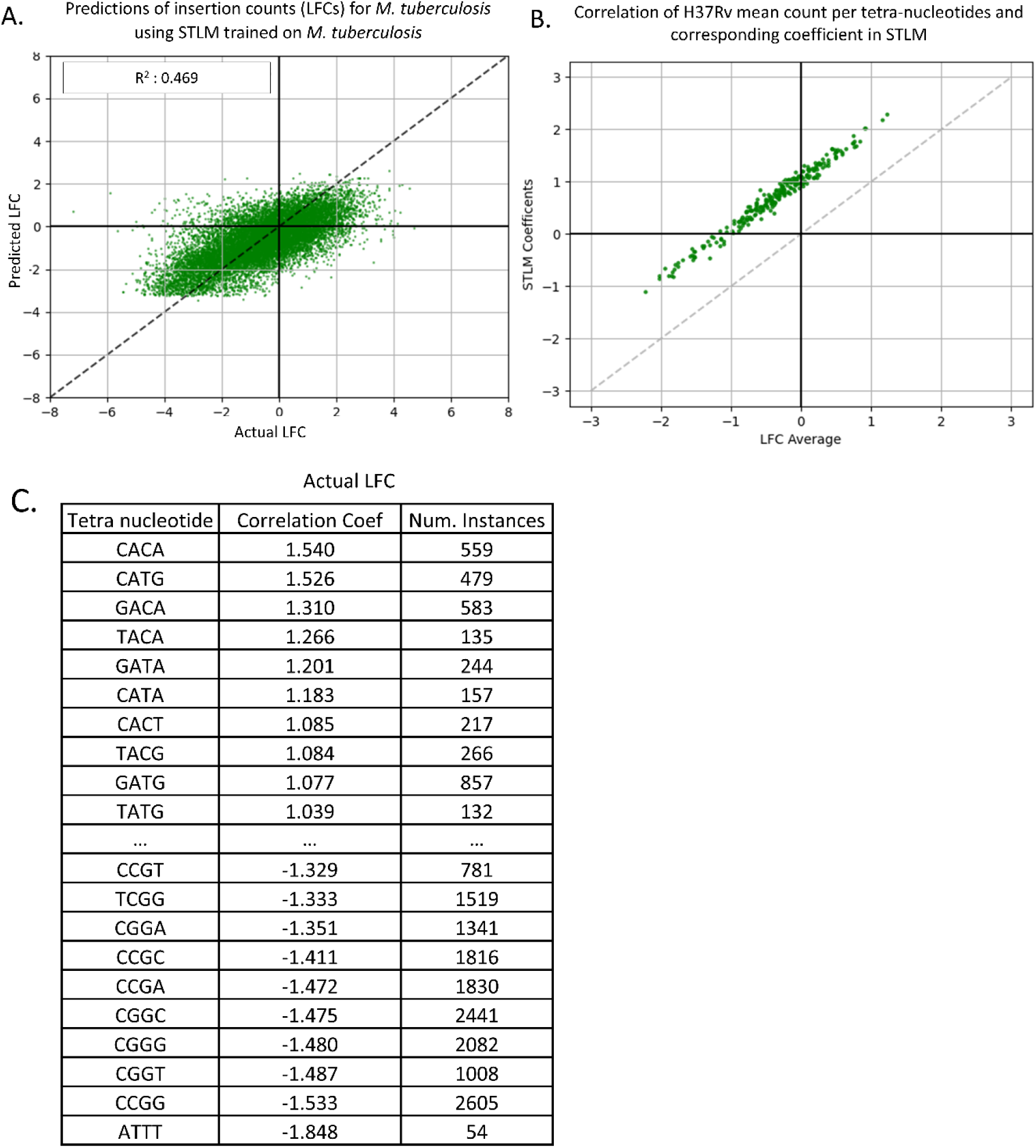
Predicted LFC vs. Actual LFC using STLM. **Panel A** shows a plot of the actual LFCs vs. the LFCs predicted by our model. The predictive power of this model is about the same as the Neural Network (R^2=^0.468) but **Panel B** shows there is a high correlation of mean LFCs of each tetra-nucleotide and the coefficient in the STLM of the same tetra-nucleotide, indicating our model represents our data well. **Panel C** shows the coefficients associated with each tetra-nucleotide (Supplemental Table T4), sorted by coefficient value.

**Figure 6:**
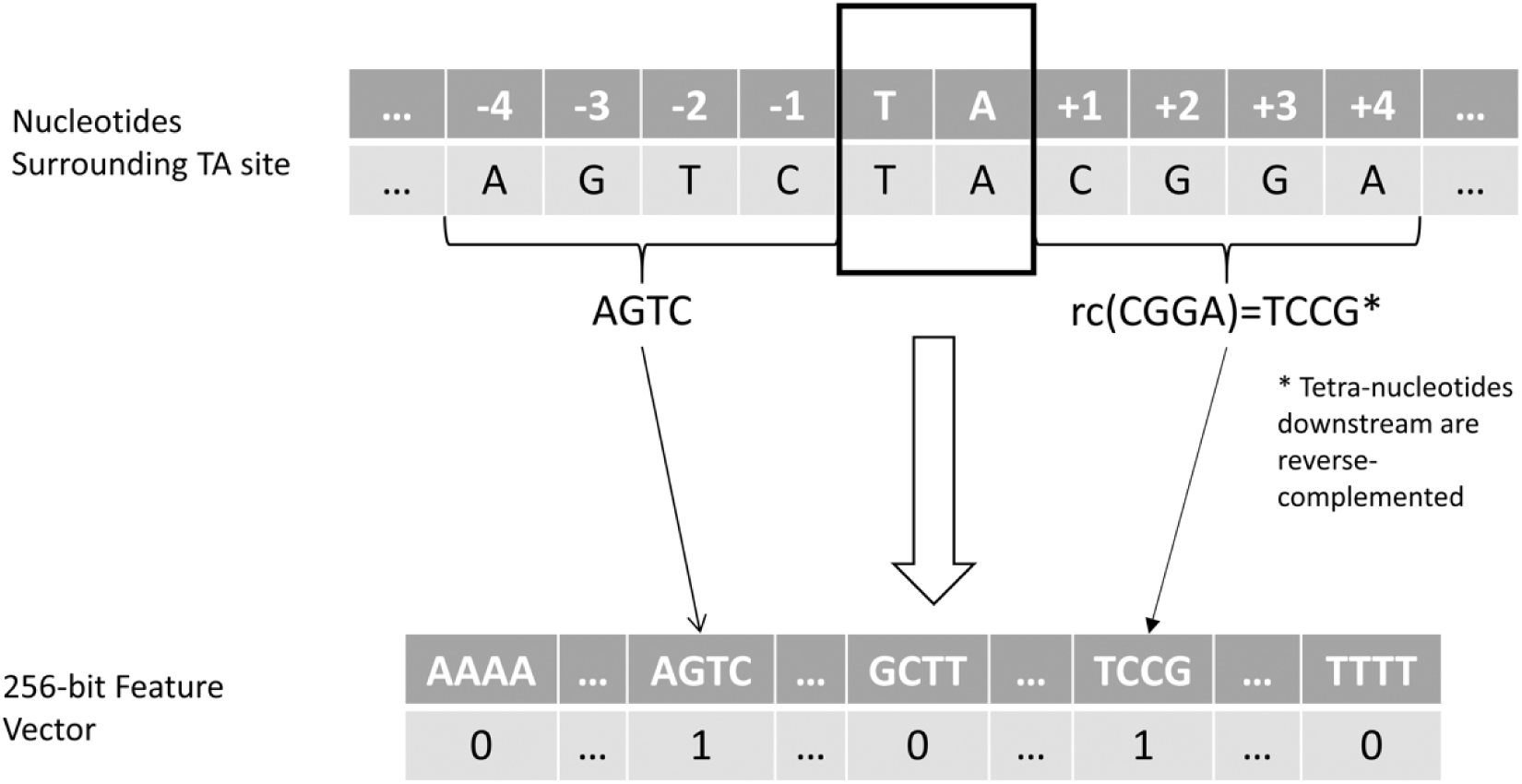
Illustration of the STLM. For each TA site the upstream tetra-nucleotide and reverse complemented (rc) downstream tetra-nucleotide are extracted. The relative bits are set in a 256-bit vector that is given as an input to the STLM to predict LFCs.

While the STLM was able to partially predict the frequency of insertion at different TA sites (R^2^=0.47), a significant amount of variability remains between observed and predicted insertion counts. This can be attributed to stochastic variability in the insertion counts across the libraries, as well other factors that the model did not account for, such as GC content outside of the -4…+4 region and DNA bendability (Lampe, Grant et al. 1998). However, when the STLM was augmented with the addition of GC content as a feature, where GC content was calculated with the ±20 bp window, it only showed an improvement in R^2^ of 0.02. When the STLM was augmented with bendability as an additional feature, calculated for each TA site using the *bend-it* program (Goodsell and Dickerson 1994), the results were nearly identical to that of the model with only the 256-bit vectors (R^2^=0.47). These experiments indicated that the tetra-nucleotides are a larger factor in the prediction of LFCs than GC content or bendability. Using only the GC content and bendability as two features to a linear model resulted in a R^2^ of nearly zero for all the datasets tested. Furthermore, plots of LFC vs. bendability and LFC vs. GC content showed little to no correlation.

### Application of STLM to other Himar1 TnSeq datasets

To evaluate whether the nucleotide biases derived from these 14 independent datasets in H37Rv are representative of generalized insertion preferences of the Himar1 transposon, we compared the biases seen so far to those in other Himar1 TnSeq datasets.

Staying within the *Mycobacterium* genus, we obtained datasets from Himar1 TnSeq libraries, grown in regular growth medium (7H9), of *M. avium* (Dragset, Ioerger et al. 2019), *M. abscessus* ATCC 19977 (Akusobi et.al, https://www.biorxiv.org/content/10.1101/2021.07.01.450732v1)*, M. smegmatis* mc^2^ 155 ΔLepA (unpublished data, E.J. Rubin), and *M. tuberculosis* H37Rv ΔRv0060 (Zaveri, Wang et al. 2020). We extracted the LFCs from the datasets based on the insertion counts at TA sites in each genome along with tetra-nucleotide vectors based on the nucleotides surrounding each TA site. The heatmaps for each of the datasets in Figure 7 shows the mean LFCs associated with each nucleotide at each position within a ±20bp window of the TA Site. These heatmaps look nearly identical to the heatmap of H37Rv in Figure 4. They exhibit the same negative LFC bias for -3 ‘C’, +3 ‘G’, -2 ‘G’, +2 ‘G’, and the same -3 ‘A’, +3 ‘T’ positive LFC bias. STLM LFC predictions for each of the new Tn-Seq datasets were adjusted by a simple regression-based procedure to correct for differences in the LFC distribution (further described in Methods). Results, calculated as correlations between predicted and observed LFCs with the regression adjustment (see Supplemental Figure S6), along with the nucleotide biases observed in the heatmaps, show that the STLM can help explain the variability in insertion counts at different TA sites for these datasets (using coefficients trained on *M. tuberculosis* H37Rv data but applied to datasets from other mycobacterial species). The predictive power of our model on the *M. abscessus* dataset (R^2^ of 0.504) is slightly higher than, but about the same as, the Mtb test set. Thus, we can explain ∼50% of the variance in insertion counts in this dataset based on the nucleotide biases. The predictive power of our model on some datasets, such as *M. avium* was lower (R^2^ of 0.262), but they still exhibited a correlation between observed and predicted LFCs (and hence insertion counts) and displayed a nucleotide pattern similar to the heatmaps from the other mycobacterial TnSeq datasets.

**Figure 7:**
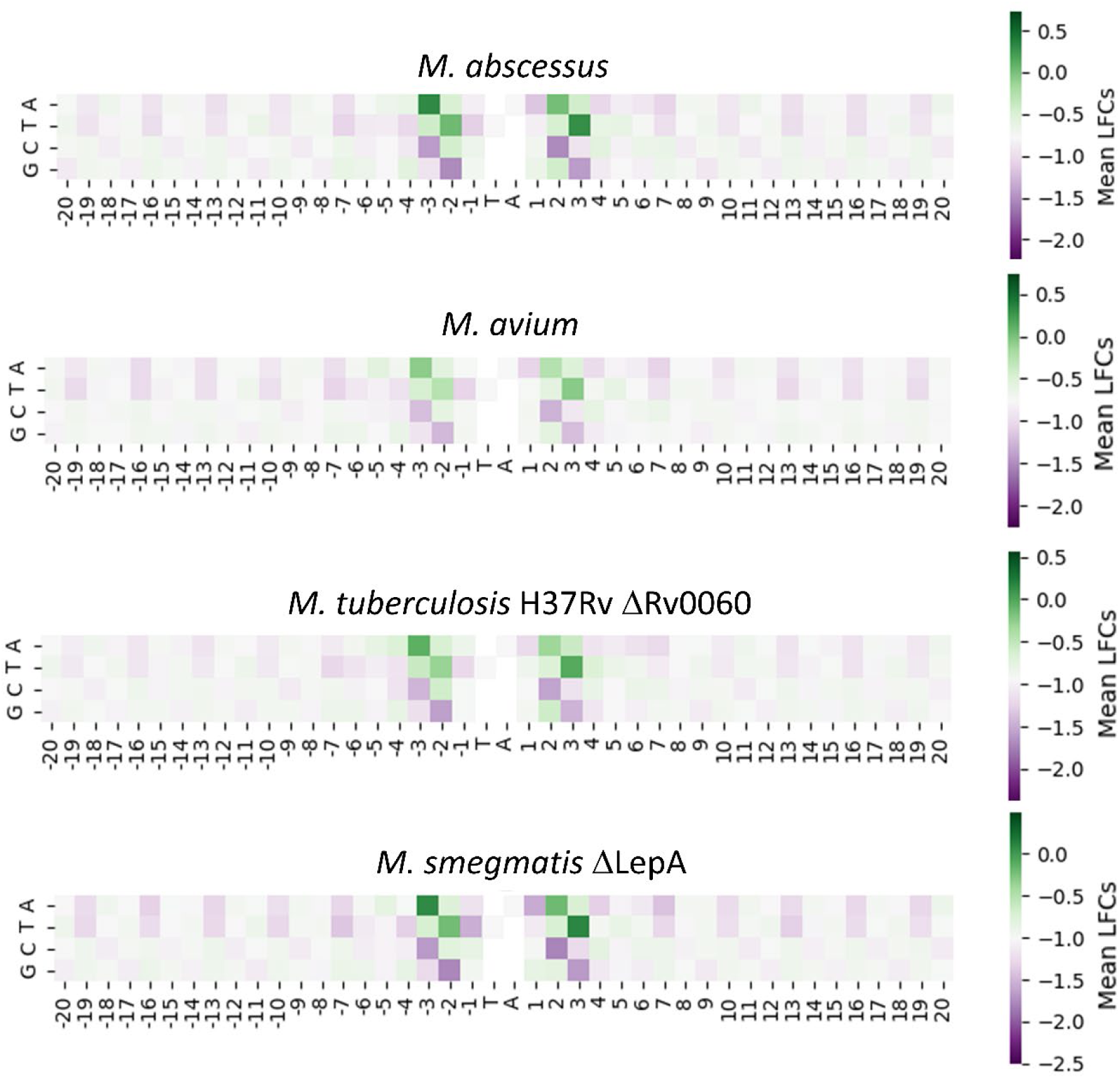
Enrichment and Depletion surrounding TA sites for Mycobacterial Tn-Seq Datasets. The four heatmaps are calculated in the same manner that the H37Rv heatmap was calculated in Figure 4. The mean of each filtered nucleotide at every position in a ±20 bp window around the TA sites is calculated. The patterns of all the heatmaps look very similar to both each other and to the H37Rv heatmap in Figure 4.

To examine whether these biases also occur outside of the *Mycobacterium* genus, we obtained Himar1 TnSeq datasets from *Caulobacter crescentus* (Murray, Panis et al. 2013)*, Rhizobium leguminosarum* (Perry, Akter et al. 2016), and *Vibrio cholera* (Chromosome I only; Chromosome II behaves similarly) (Chao, Pritchard et al. 2013). We calculated LFCs (log-fold-change of insertion counts relative to local mean) at each TA site in these genomes and plotted the heatmaps as associations of nucleotides at specific positions around the TA with LFCs. As Figure 8 shows, the heatmaps associated with all three datasets reflect the same nucleotide patterns found in the mycobacterial datasets. Applying the STLM to these datasets yielded significant correlations between predicted and observed LFCs, with statistically significant R^2^ values (see Supplemental Figure S7). The correlation for *Vibrio cholera* is lower than the others (R^2^= 0.249) possibly due to sequence preferences in the fragmentase used for shearing during the sample prep for sequencing. This was done differently than other TnSeq experiments and could have introduced additional variance into the insertion counts for the *Vibrio* dataset. However, the heatmap shows a pattern consistent with the nucleotide biases we see with the Tn-Seq datasets from other organisms. This indicates that the nucleotide biases visible in the mycobacterial datasets also explains some of the insertion count variances present in non-mycobacterial datasets, thus supporting that the STLM captures generalized site-specific biases on insertion preferences of the Himar1 transposon.

**Figure 8:**
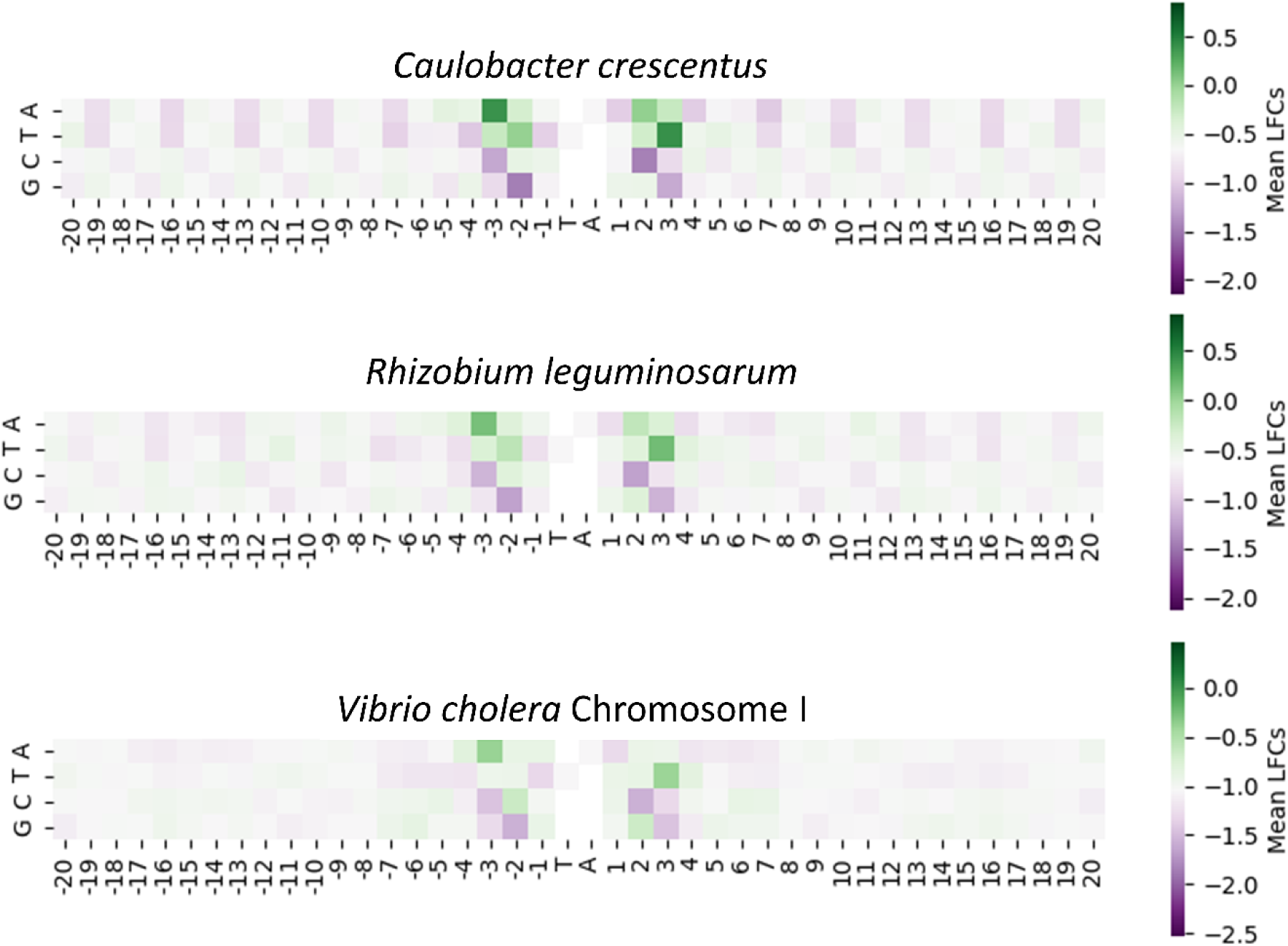
Enrichment and Depletion surrounding TA sites for non-Mycobacterial Tn-Seq Datasets. The four heatmaps are calculated in the same manner that the H37Rv heatmap (Fig. 4) and mycobacterial heatmaps (Fig. 7) were calculated. The mean of each filtered nucleotide at every position in a ±20 bp window around each TA site is calculated, and centered on the median of mean LFCs.

### SNPs around TA sites in an *M. abscessus* Clinical Isolate Exhibit Predictable Changes in Insertion Counts

To evaluate whether changes in nucleotides proximal to TA sites would have a predictable effect on transposon insertion counts, we obtained a Himar1 Tn library for a clinical isolate of *M. abscessus* Taiwan49 (*Mab* T49) and compared it to a Tn library in the reference strain, ATCC 19977 (generated by methods described in the accompanying manuscript by Akusobi et al. https://www.biorxiv.org/content/10.1101/2021.07.01.450732v1; see Availability for data files with raw insertion counts). These two strains of *M. abscessus* are fairly divergent, belonging to different subspecies (ATCC 19977 in *Mab subsp. abscessus*, and Taiwan49 in *Mab subsp. massiliense*); they have 114,335 SNPs between them based on a genome-wide alignment. However, at the level of functional genomics, they are similar. As determined through the HMM method in TRANSIT (DeJesus, Ambadipudi et al. 2015), 513 out 4923 total genes in ATCC 19977 and 451 out 4225 total genes in T49 are predicted to be essential or growth defect genes. 417 of these genes overlap. Figure 9 shows that predicting insertion counts in this isolate with the STLM yielded an R^2^ value of 0.49. This is, as expected, quite similar to the results of the *M. abscessus* reference dataset reported above. After aligning the genomes *M. abscessus* Taiwan49 clinical isolate and *M. abscessus* reference strain, we found 9303 TA sites where there was exactly one SNP in the 8-nucleotide window (±4 bp) surrounding the TA site.

**Figure 9:**
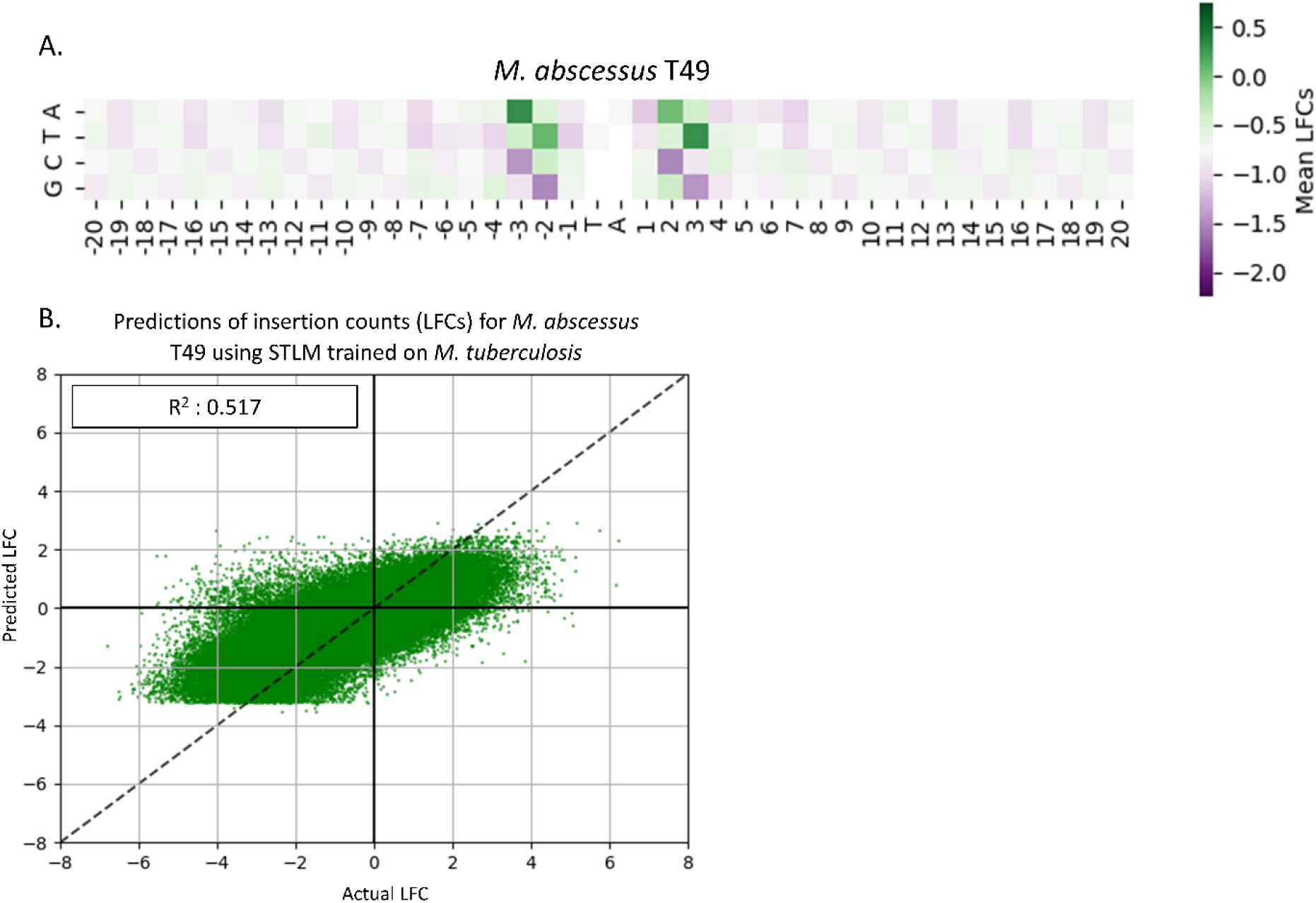
*M. abscessus* Taiwan49 Clinical Isolate Dataset. The heatmap in **Panel A** is calculated in the same manner that the previous heatmaps were calculated. The pattern of this heatmap looks very similar to the H37Rv heatmap (Fig. 4) as well as the heatmap for the *M. abscessus* ATCC 19977 reference strain (Fig. 8). The predictive power of the STLM on the Mab T49 dataset in **Panel B** shows a high R^2^ value of 0.517, like that of the *M. abscessus* reference dataset. This indicates that nucleotide biases explain at least half of the variance in insertion counts for this dataset with nucleotide biases.

A plot of the average changes in observed LFCs versus the average changes in predicted LFCs between the reference and isolate strain at these sites for every TA site with an adjacent SNP can be seen in Figure 10 (see Methods and Materials). We expected that when a nucleotide with a high negative bias was mutated, the observed LFC would increase, and when a nucleotide with a high positive bias was changed, the observed LFC would decrease. Figure 10 shows this effect. The colored points in the graph are the most significant nucleotide-position pairs that we have previously observed to have the highest LFC biases. When a nucleotide is switched from an ‘A’ in -3 position (blue) or a ‘T’ in the +3 position (green) to any other nucleotide, there is a decrease in observed LFC and when a ‘G’ in -2 position (orange) or ‘C’ in +2 position (pink) is changed, there is an observed increase. The presence of this effect of SNPs on the LFC i.e., differences in insertion counts at corresponding TA sites in different clinical isolates, along with the high correlation of the observed and predicted LFC changes, provides further evidence that the STLM can represent the nucleotide biases on transposon insertion preferences with high accuracy.

**Figure 10:**
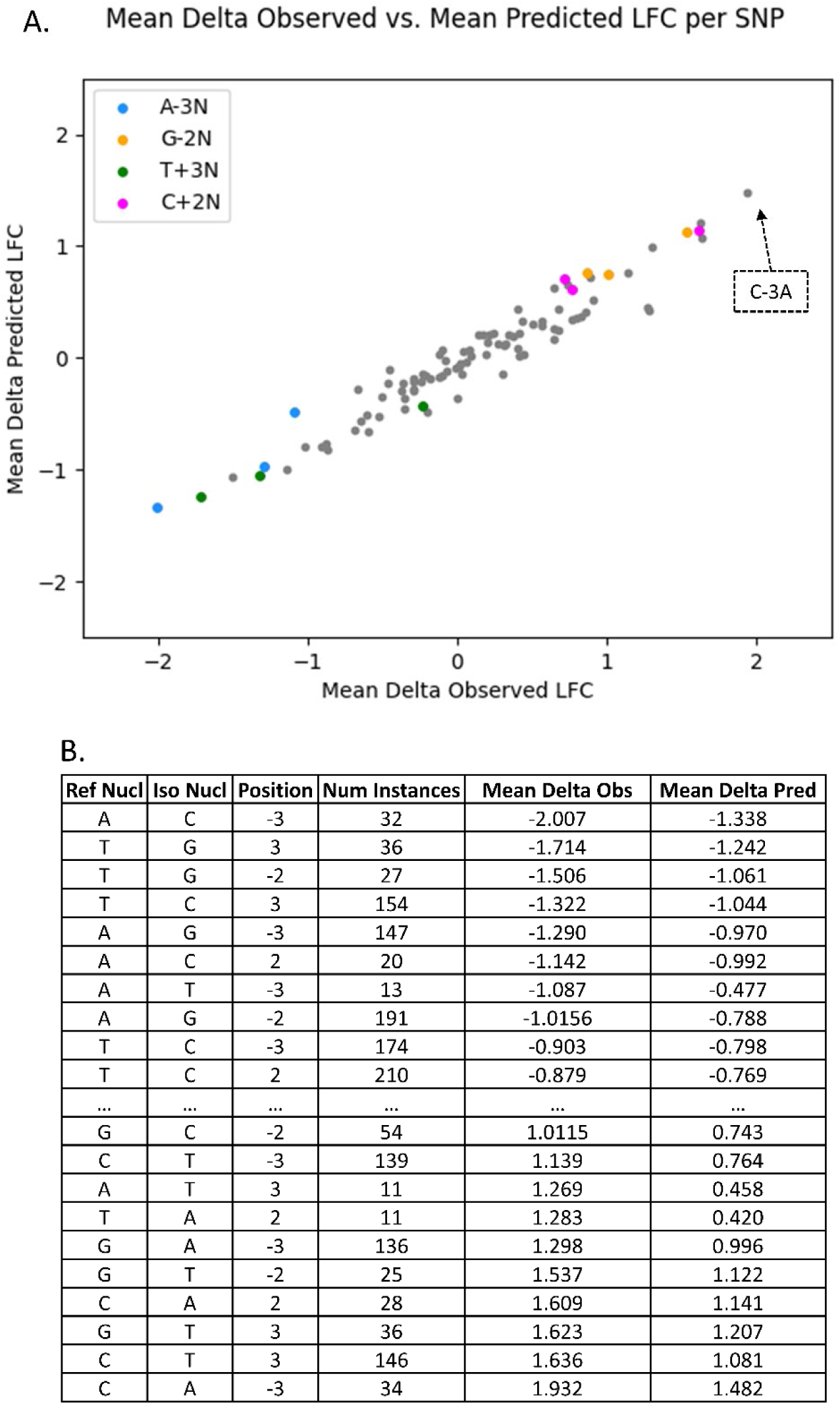
SNPs in *M. abscessus* Taiwan49 Clinical Isolate Exhibit Predictable Changes in Nucleotide Biases. **Panel A** shows the correlation of changes in observed vs predicted LFCs for the 96 possible SNPs in the - 4…+4 window from the TA site. The colored markers are the nucleotide-position pairs previously found to have the highest biases. The table in **Panel B** is sorted by increasing mean delta observed LFC, provides more details on these SNPs. As expected, the most extreme changes occur when the SNP occurs in the -3, -2, +2, or +3 positions. The top 10 and bottom 10 values i.e., the biggest decreases and biggest increases in LFC follow the heatmap patterns of the Himar1 datasets tested.

The accompanying table in Figure 10 is a truncated view of the SNPs sorted in increasing order of mean observed LFC change (for full table see Supplemental Table T2). In addition to the general pattern observed in the plot, we see that the magnitudes of the LFC differences correspond to the magnitudes of nucleotide biases. In the previous heatmaps, ‘A’ in the -3 position (and the downstream reverse complemented pattern) shows the strongest bias for high LFCs and ‘C’ in the -3 position or ‘G’ in -2 position (and the downstream reverse complemented pattern) shows the strongest bias towards low LFCs. Following this pattern, the biggest decrease in mean observed LFC, occurred when an ‘A’ in the -3 position was changed to a ‘C’ and the biggest increase in mean observed LFCs occurred when ‘C’ in the -3 position was changed to an ‘A’. Thus, the effect of SNPs between a pair of moderately divergent strains correspond to the nucleotide biases observed within various Himar1 datasets and furthers the notion that these biases are general and can explain a significant portion of the variance in insertion counts of Himar1 Tn-Seq datasets.

### Using Expected Insertion Counts to Improve Gene Essentiality Predictions

Previous methods of identifying essential genes within individual datasets have been based on the magnitude of insertion counts. For example, tools such as TnSeq-Explorer (Solaimanpour, Sarmiento et al. 2015) use the mean of insertion counts in sliding windows to classify genes by essentiality. The limitation of relying on raw insertion counts is that they can be highly variable among TA sites, and this noise can lead to inaccurate estimation of the relative level of fitness defects caused by transposon disruption. We describe a new method, called the TTN-Fitness method using the Gene+TTN model, which considers the site-specific biases on Himar1 insertion preferences to correct the observed counts for expectations based on the nucleotides surrounding each site.

The Gene+TTN model incorporates nucleotide context into an insertion count based model, allowing us to decouple the two main causes for low insertion counts: biological, and Himar1 insertion preferences. This allows us to make a more informed assessment on the level of gene fitness defect for biological reasons. The input to the model for each TA site is a vector consisting of a binary encoding of the gene in which it is located, combined with the 256 tetra-nucleotide (TTN) features. Each TA site is represented as a bit vector, with 3981 features, one for each gene, and 256 features encoding the upstream and reverse-complemented downstream tetra-nucleotides adjacent to the site. We excluded TA sites from genes determined to be ‘Essential’ through the Gumbel analysis and Bernoulli Distribution. The model can be represented in matrix form as:

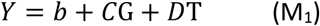

where Y is a vector of the log10 of insertion counts at every TA site, G is the matrix of 3981 gene covariates for each site, C is vector coefficients to be fit per gene, T is the matrix of 256 tetra-nucleotide covariates for each site and D is the vector of coefficients to be fit per tetra-nucleotide. The intercept b is close to the global average of log10 insertion counts and the coefficients (C) for every gene reflect the deviation of the gene’s mean log10 insertion count from the global average, adjusting for the effect of surrounding nucleotides (D). Essentially, we are finding the deviation of the gene’s mean insertion count from the global average based on biological reasons, i.e., subtracting out the effect of site-specific nucleotide based Himar1 insertion preferences. Thus, the gene-specific coefficients (C) represent adjusted estimates of the fitness level of each gene.

The regression model was trained on the *Mtb* H37Rv in-vitro dataset. The significance of genes (i.e. p-value) was calculated using a Wald test (Draper and Smith 1998), and then adjusted for multiple testing to limit the False Discovery Rate (FDR) to ≤5% using the Benjamini-Hochberg method (Reiner, Yekutieli et al. 2003). Genes with an adjusted p-value < 0.05 and negative coefficient are interpreted as ‘Growth Defect’ (GD) genes, and those with adjusted p-value < 0.05 and positive coefficient are interpreted as ‘Growth Advantaged’ (GA) genes. Genes with an insignificant coefficient near 0 (adjusted p-value > 0.05) are interpreted as ‘Non-Essential’ (NE). Genes identified a priori as essential by the Gumbel method in TRANSIT (DeJesus, Ambadipudi et al. 2015) were marked ‘Essential’ (ES) by the TTN-Fitness method and excluded from both training and testing. Gumbel identifies large essential genes well but tends to classify small genes (with <10 TA sites) as ‘Uncertain’, depending on the overall level of saturation of the dataset. Thus, we use the Bernoulli distribution to classify additional significant genes (p<0.05) lacking insertions that are likely essential as ‘Essential-B’ (ESB, as a subcategory of ES) (see Methods and Materials). The HMM+NP model, a modified HMM to account for non-permissive sites described by DeJesus (DeJesus, Gerrick et al. 2017), distinguishes between ‘ES’ and ‘ESD’ (Domain-Essential) genes, which our model does not. For model comparison, we have combined the two categories into one labeled ‘ES/ESD’. As seen in Figure 11A, the TTN-Fitness method labels a similar number of genes essential as the HMM-NP method, though slightly fewer non-essential and more in the growth-defect and growth-advantaged categories (DeJesus, Gerrick et al. 2017). The confusion matrix in Figure 11B shows that there are 345 genes labeled ‘Essential’ in both the TTN-Fitness method and the HMM+NP model (i.e. on the diagonal in the confusion matrix), showing a great deal of overlap. 1777 ‘Non-Essential’ genes also overlap between the 2 methods. However, the biggest difference is that large number of genes labelled as ‘Non-Essential’ (NE) by the HMM get reclassified as either ‘GD’ or ‘GA’ by the TTN-Fitness method. 14.7% of genes labeled ‘Non-Essential’ in the HMM+NP model have slightly lower than average insertion counts and are classified as ‘GD’ via the Gene+TTN (M_1_) model. 25.4% of genes labeled ‘Non-Essential’ by the HMM+NP model have insertion counts slightly higher than average and are classified as ‘GA’ through the Gene+TTN (M_1_) model. This shows that the TTN-Fitness method labels genes similarly to the HMM+NP model for the most part, but is more sensitive to deviations from the average insertion count and consequently labels some genes more specifically as ‘GD’ or ‘GA’.

**Figure 11:**
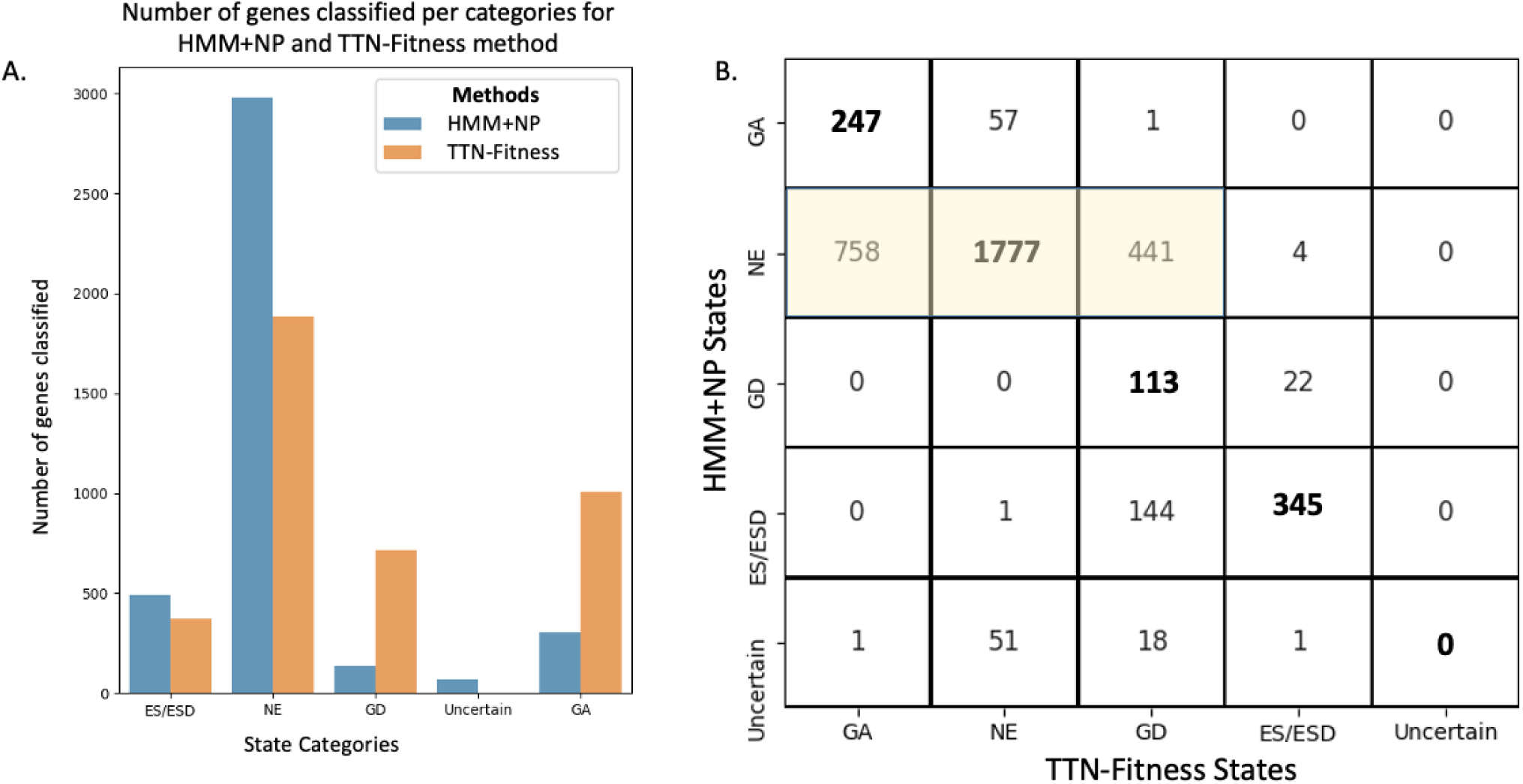
Distribution of HMM+NP and TTN-Fitness states for genes provided in the *Mtb* H37Rv dataset. **Panel A** shows the distribution of classification of genes by the two methods. **Panel B** shows the confusion matrix of the classification of genes in the two methodologies. Most of genes are labeled NE in both models. Genes determined to be “Uncertain” in the HMM+NP model are assigned other states in the TTN-Fitness method. A fraction of genes labeled “NE” in the HMM+NP model (highlighted matrix components) are reassigned to be “GA” or “GD” using the TTN-Fitness method, indicating that the TTN-Fitness method is more sensitive in estimating fitness than the HMM+NP model.

Figure 12A shows a linear relationship between the coefficients associated with tetra-nucleotides features in the Gene+TTN (M_1_) model and the corresponding coefficients of the STLM, illustrating that the influence of tetra-nucleotides on predicted counts captured in this model is consistent with the effect previously discussed in the STLM. <u>Panel B</u> shows the difference in the fitness assessment of genes when compared to a Gene-Only (M_0_) model, dropping the TTN features and hence lacking the site-specific adjustments based on tetra-nucleotide covariates. The Gene-Only (M_0_) model encodes only the gene at every TA site and can be expressed in matrix form as:

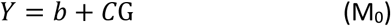

**Figure 12:**
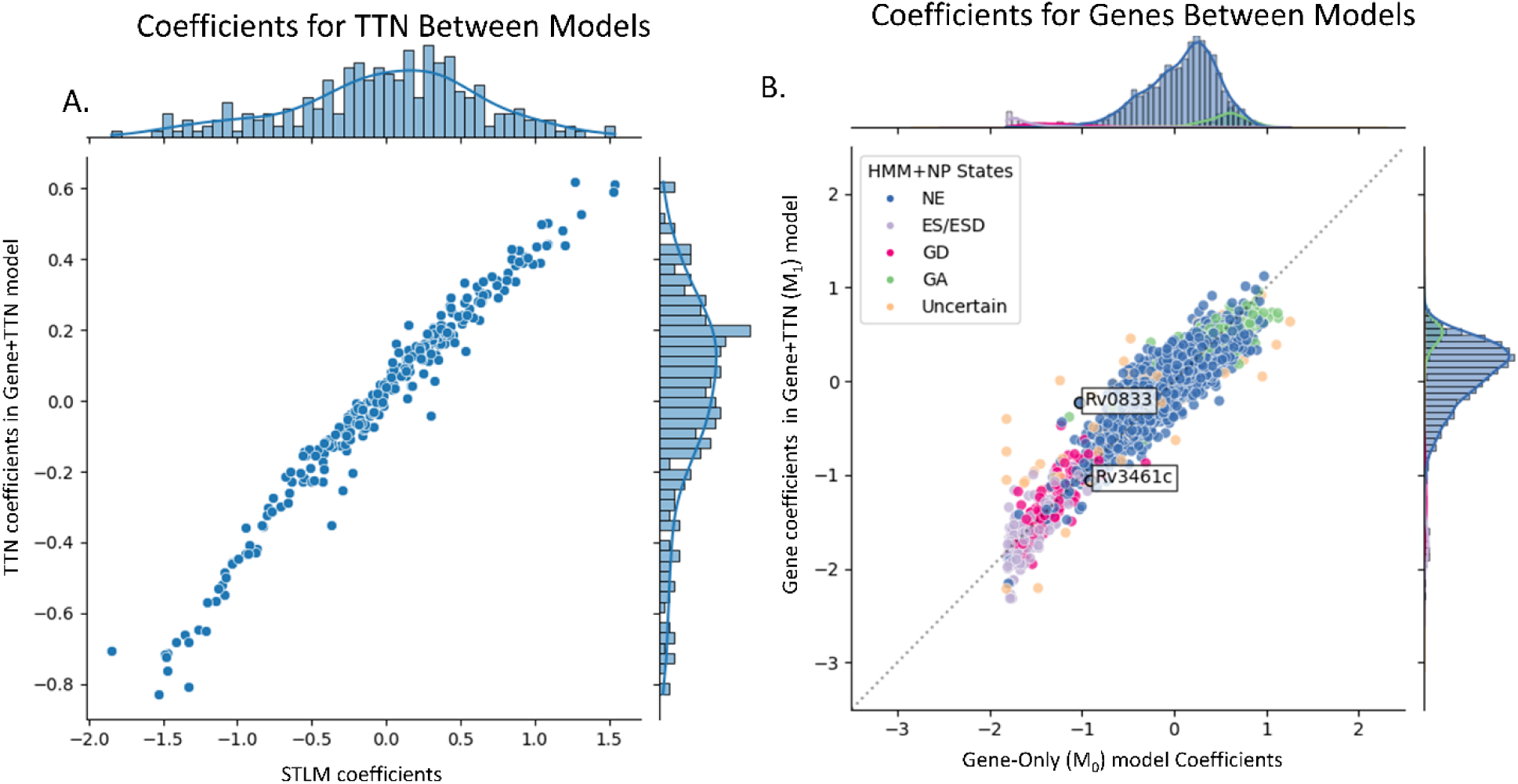
Correlation of coefficients in Gene+TTN model (of the TTN-Fitness method) and coefficients in models using its components. Correlation of gene coefficients between the Gene-Only model and the TTN Fitness model **(Panel A**) show a linear trend, indicating that most genes behave in the same way and yield similar results in both models. However, there are a few that are show log fold change greater than this majority. The scale of coefficients in the Gene+TTN model is greater than the Gene-Only model, indicating a notable number of gene’s predicted fitness estimate changes with the inclusion of nucleotide context. The points with black outlines and labels are genes that we have explored. Correlation of coefficients of TTNs in the STLM and the TTN Fitness Model **(Panel B)** has a strong linear relationship as well as similar distributions, indicating that the models incorporate in the effects of TTNs on the insertion count in the same way.

The intercept (b) is the global average log10 insertion count in the genome and the coefficient C corresponding to each gene is the deviation of the gene’s mean insertion count from the global average. As this model does not incorporate the tetra-nucleotides, if there is a gene with a very negative coefficient, it will be interpreted to be ‘Growth Defect’ regardless of whether the suppression of insertions is due to true biological gene defect or nucleotide bias. The scatterplot of the gene coefficients between the two models in <u>Panel A</u> shows a strong linear trend, indicating estimated mean (log10) insertion counts for most genes are quite similar between the two models. However, the dispersion suggests that taking the nucleotide context into account changes the fitness estimate for a number of outlying genes. Genes that show the highest differences in coefficients between the two models are frequently labeled as ‘Uncertain’ by the HMM+NP model (DeJesus, Gerrick et al. 2017), a majority of which are small genes with fewer than 5 TA sites. Details on the difference in the coefficients and their significance (determined through a Student t-test and an FDR adjusted p-value) can be found in the Supplemental Table T3.

An example of a gene whose fitness interpretation is changed via the Gene+TTN model in the TTN-Fitness method (compared to Genes-only model), to better reflect its biological significance, is Rv0833 (PE_PGRS13). The gene is seen in Figure 12B is interpreted as ‘Growth Defect’ through model M_0_ (Genes-only; C = -1.02, adjusted p-val = 2.95 x 10^−6^) and as ‘non-essential’ by model M_1_ (Gene+TTN; C = -0.26, adjusted p-value = 0.109, hence not significantly different from 0). The difference in labeling indicates that, based on the surrounding nucleotides, the low insertion counts at TA sites in Rv0833 are expected. This is supported by the fact that the PE_PGRS genes are especially GC-rich (Gey van Pittius, Sampson et al. 2006). The gene contains 12 TA sites spanning 2250 base pairs. 81.3% of the nucleotides were ‘G’s or ‘C’s and 6 sites contained the non-permissiveness pattern. Thus, observed insertion counts in the gene are much lower than the global average insertion count, but they are expected to be. Although studies suggest that genes within the PE/PPE family may be involved in inhibition of antigen processing in hosts, PE_PGRS genes have been shown to be non-essential in-vitro (Gey van Pittius, Sampson et al. 2006). Thus, the Gene+TTN model was able to evaluate the fitness of PE_PGRS13 more accurately than the Gene-Only model, demonstrating that incorporating the nucleotide context surrounding each TA site improves the fitness assessment of this gene.

To investigate genes that exhibit large differences in fitness assessment between the TTN- Fitness method and the HMM+NP method, Figure 13 shows a volcano plot of the gene coefficients from the Gene+TTN model versus the –log_10_ of the FDR-adjusted p-value. The gray points in the plot are gene coefficients that were not seen to significantly deviate from 0. These are interpreted as ‘non-essential’ genes by the TTN-Fitness method. The genes that were found to be significant are colored according to their labels in the HMM+NP model. The vertical solid line at C=0 is where the colored genes on the left are interpreted as ‘GD’ and colored genes on the right are interpreted as ‘GA’ by the TTN-Fitness method. All significant genes labeled ‘GA’ or ‘GD’ by the HMM+NP model fall on their respective sides of the C=0 line, but there are a few ‘non-essential’ and ‘Uncertain’ genes that are reclassified by the TTN-Fitness method.

**Figure 13:**
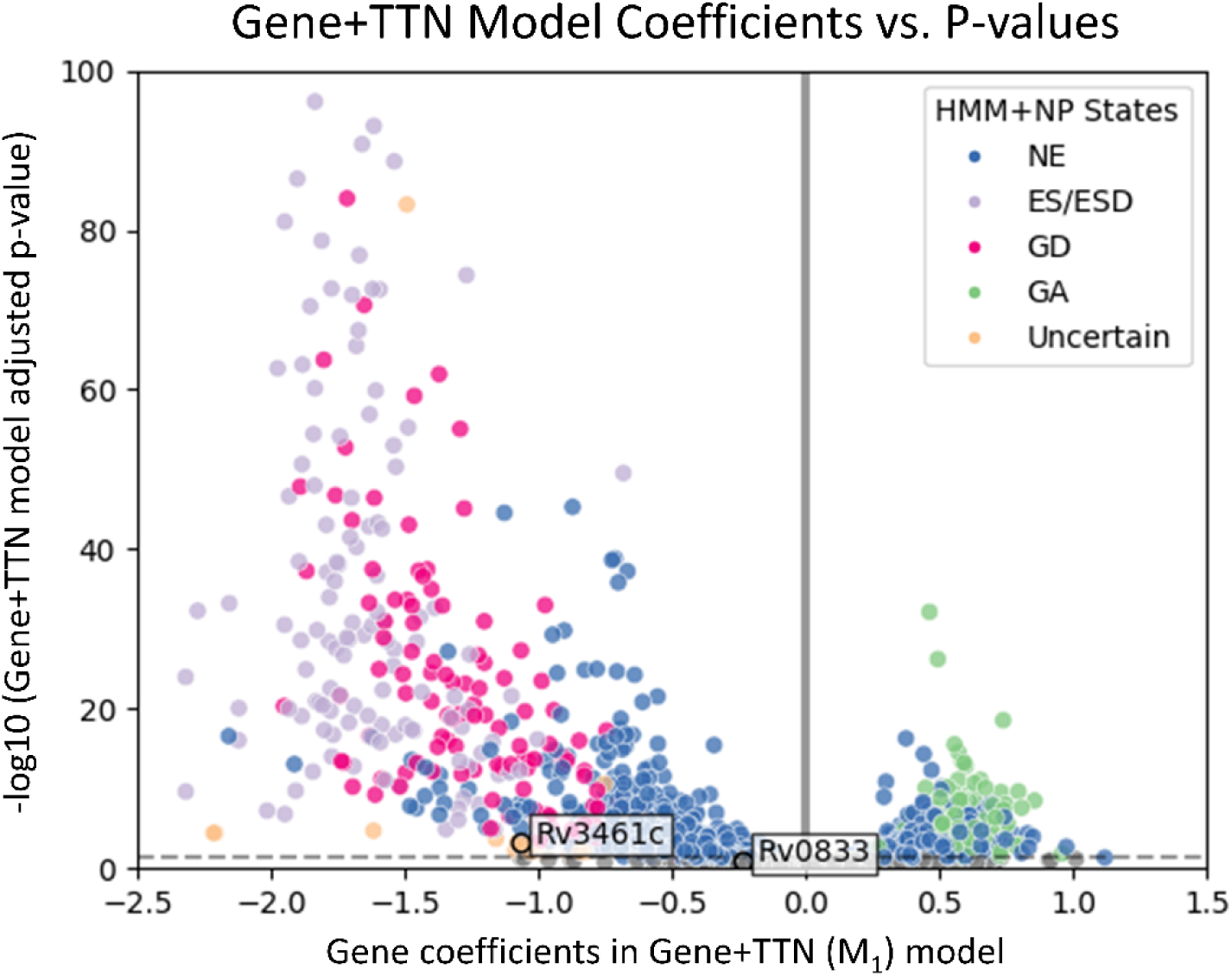
Plots of gene coefficients versus adjusted p-values in Gene+TTN model, colored by states determined by the HMM+NP model. The HMM+NP methodology labels genes as “Non-Essential” (NE), “Essential” (ES/ESD), “Growth Defect” (GD), “Growth Advantage” (GA) and “Uncertain”. “Uncertain” genes are typically smaller genes. The horizontal dashed line is where adjusted p-value = 0.05 in the Gene+TTN model. By the TTN-Fitness method, genes below that line are insignificant (gray) and thus “NE”. The vertical solid line is where gene coefficient C=0 in the Gene+TTN model. By the TTN-Fitness method, colored points left of the line are “GD” genes and colored points to the right are “GA” genes. The genes with labels are discussed in the text.

With improvements in fitness assessment from the incorporation of tetra-nucleotides, small genes (3 or less TA sites) labeled “Uncertain” by the HMM+NP model can be evaluated with greater confidence. Of the 71 genes labeled “Uncertain” by the HMM+NP model, most (65) have 3 or fewer TA sites, indicating the uncertainty comes from the short length of the gene. These genes are all concretely classified by the TTN-Fitness method (mostly as ‘Non-Essential’ (51) or ‘Growth-Defect’ (18); see Figure 11B) .Rv3461c (*rpmJ*, 50S ribosomal protein L36), a gene with 3 TA sites, is an example of such an “Uncertain” gene (DeJesus, Gerrick et al. 2017). The gene is seen in Figure 12B to be interpreted as ‘Non Essential’ by the Gene-Only model M_0_ (C = -0.87, adjusted p-val = 0.074, not significantly different from 0) and ‘Growth Defect’ by the Gene+TTN (M_1_) model(C = -1.02, adjusted p-val = 9.41 × 10^−4^), indicating the insertions for the genes are lower than expected according to the surrounding tetra-nucleotides. Figure 13 shows that the gene is similar to other genes labeled ‘Growth defect’ or ‘Essential’ by the HMM+NP model. Rv3461c is a part of the L3P family of ribosomal proteins. Other genes in this family have been labeled as ‘Essential’ or ‘Growth Defect’ by the HMM+NP model and ‘Growth Defect’ per the Gene+TTN model. In fact, *rpmJ* was categorized as a ‘Growth Defect’ gene in early TraSH experiments (Sassetti, Boyd et al. 2003). Therefore, this previously ‘Uncertain’ gene should be interpreted as ‘Growth Defect’ (possibly even ‘Essential’), as the TTN-Fitness method suggests, with confidence.

These examples show the improvement of fitness assessment with the incorporation of tetra-nucleotides in an insertion-count only model. This enables the TTN-Fitness method to account for the effect of genomic context on the Himar1 transposon insertion preferences, and thus better assess a gene’s fitness defect due to genuine biological causes.

## Discussion

Previous studies have demonstrated the presence of some site-specific biases on Himar1 transposon insertion preferences based on a non-permissive pattern that exists around TA sites with low insertion counts in non-essential regions (DeJesus, Gerrick et al. 2017). This led us to hypothesize that perhaps insertion counts at different TA sites could be predicted based on surrounding nucleotides. We developed a model that captures nucleotide biases and uses them to predict changes in relative insertion counts i.e., LFCs. The LFC metric compares raw counts at a site to the local average, which allows us to predict the deviation in insertion counts from the neighborhood rather than the absolute insertion counts themselves. This method allows us to examine just the effect of the nucleotides on the insertion counts, independent of biological effects (e.g., genes with different levels of growth defect). The STLM developed for the task incorporated tetra-nucleotides upstream and downstream of the TA site, taking advantage of the symmetric nature of the bias patterns observed in the heatmaps. Furthermore, the tetra-nucleotide features ensured that the model could capture non-linear combinations (interactions) of nucleotides proximal to the TA site, not just incorporating the effects or individual nucleotides in an additive way. The STLM statistically performed as well as the neural network, and in addition was able to provide further insight into nucleotide patterns that influence insertion counts.

The coefficients of the trained STLM showed that there was a pattern of insertion count suppression consistent with the non-permissive pattern previously observed (DeJesus, Gerrick et al. 2017). In addition, a pattern of increased insertion counts in the presence ‘A’ in the -3 position or ‘T’ in the +3 position was also visible. But the linear model represents these patterns in a more general way so that they can be used to predict expected insertion counts at any TA site, conditioned on the surrounding nucleotides. These nucleotide biases were able to explain up to ∼50% of insertion count variance in the other Himar1 datasets. These site-specific nucleotide biases were observed in a variety of TnSeq datasets from other mycobacterial and non-mycobacterial species. Comparing TA sites with substitutions in the ±4 bp window between two divergent strains of *M. abscessus* showed changes in observed LFCs that corresponded to nearby SNPs as predicted by the STLM, providing further evidence of the generality of these biases.

There is a precedent for transposons in some families having insertion biases for certain sequence patterns. For example, even though the Tn5 transposase can insert anywhere in a genome, it tends to insert in GC-rich regions (Goryshin and Reznikoff 1998) (Green, Bouchier et al. 2012). Furthermore, a detailed pattern analysis applied to known Tn5 insertion sites suggested that the consensus pattern for preferred target sites is A-GNTYWRANC-T (Goryshin, Miller et al. 1998). Tc1 (also in *mariner* family) was shown to weakly prefer inserting at TA sites with this consensus pattern: CAYATATRTG (Korswagen, Durbin et al. 1996). The pattern included a coupled symmetric target site preference of an ‘A’ in position -3 and a ‘T’ in position +3, consistent with our model. We were able to identify similar sequence-dependent patterns and quantify them in a more general way with a model that can predict expected insertion counts for every TA site.

Early studies in *E. coli* suggested that the Himar1 transposon tends to insert at TA sites in more “bendable” regions of the genome (Lampe, Grant et al. 1998), as measured experimentally. Bendability is a cumulative effect of specific nucleotides on local geometric parameters of the DNA helical axis; each nucleotide makes a small contribution, on the order of a few degrees, to angular distortion (bend, roll, tilt) of the axis, with different nucleotides (or combinations of nucleotides) having a different effect. This can accumulate over tens of nucleotides to produce a macroscopic bend or kink in the DNA. Goodsell and Dickerson (Goodsell and Dickerson 1994) parameterized the geometric effects for each trinucleotide and used this to generate a model which can be used to predict the bend and twist of the helical axis accumulated locally using a sliding window. It was speculated that local bendability could facilitate the melting of the double-helix, recognition/binding of the transposase, and formation of the pre-cleavage complex (Lampe, Grant et al. 1998). However, while it is possible that bendability contributes weakly to Himar1 insertion preferences, the effect likely spans a larger window of nucleotides than just ±4 bp around the TA sites; local bendability is not likely to be substantially affected by the 4 nucleotides on either side of the TA sites, which have a predominant influence according to our statistical analysis. In addition, we computed this around the TA sites in our dataset and added it as a covariate in our linear models, but it did not improve the performance of the models.

The patterns of nucleotide biases on Himar1 transposon insertion preferences may have emerged as a result of the physical interaction between the Himar-1 transposase and the DNA. Figure 14 displays the X-ray crystal structure of the complex between the Mos1 transposon (also in the *mariner* family) and the pre-cleavage state of the DNA double helix (Dornan, Grey et al. 2015). As expected, the components of the TA dinucleotide (T57, A58) interact with the protein (residues 119-124 (WVPHEL)-orange). However, the 4 adjacent nucleotides also make extensive contact with the protein in a small tunnel by packing against Asp284-His293 (green). Arg118 likely makes charged-polar interactions with the nucleotides at position -2 and -3. These positions are where different nucleotides proximal to TA dinucleotides are observed to have insertion biases in Himar1 datasets. The interactions between these TA-adjacent nucleotides and amino acid side chains in the transposase could influence the energetics and therefore the frequency of successful transposon reactions at TA sites. While it would be tempting to try to perform a detailed analysis of the hydrogen-bonding and other molecular interactions between nucleotides in the DNA fragment and amino acid side-chains of the transposase they contact to derive a structural explanation for the observed preferences for certain nucleotides surrounding the TA site, it must be remembered that this structure is of MosI (whose insertion biases are unknown, except for TA restriction), and a detailed analysis of molecular interactions relevant to the biases of the Himar1 transposase, as we have characterized, will have to await determination of an X-ray crystal structure of a complex of the Himar1 transposase bound to a target DNA fragment (containing a TA site).

**Figure 14:**
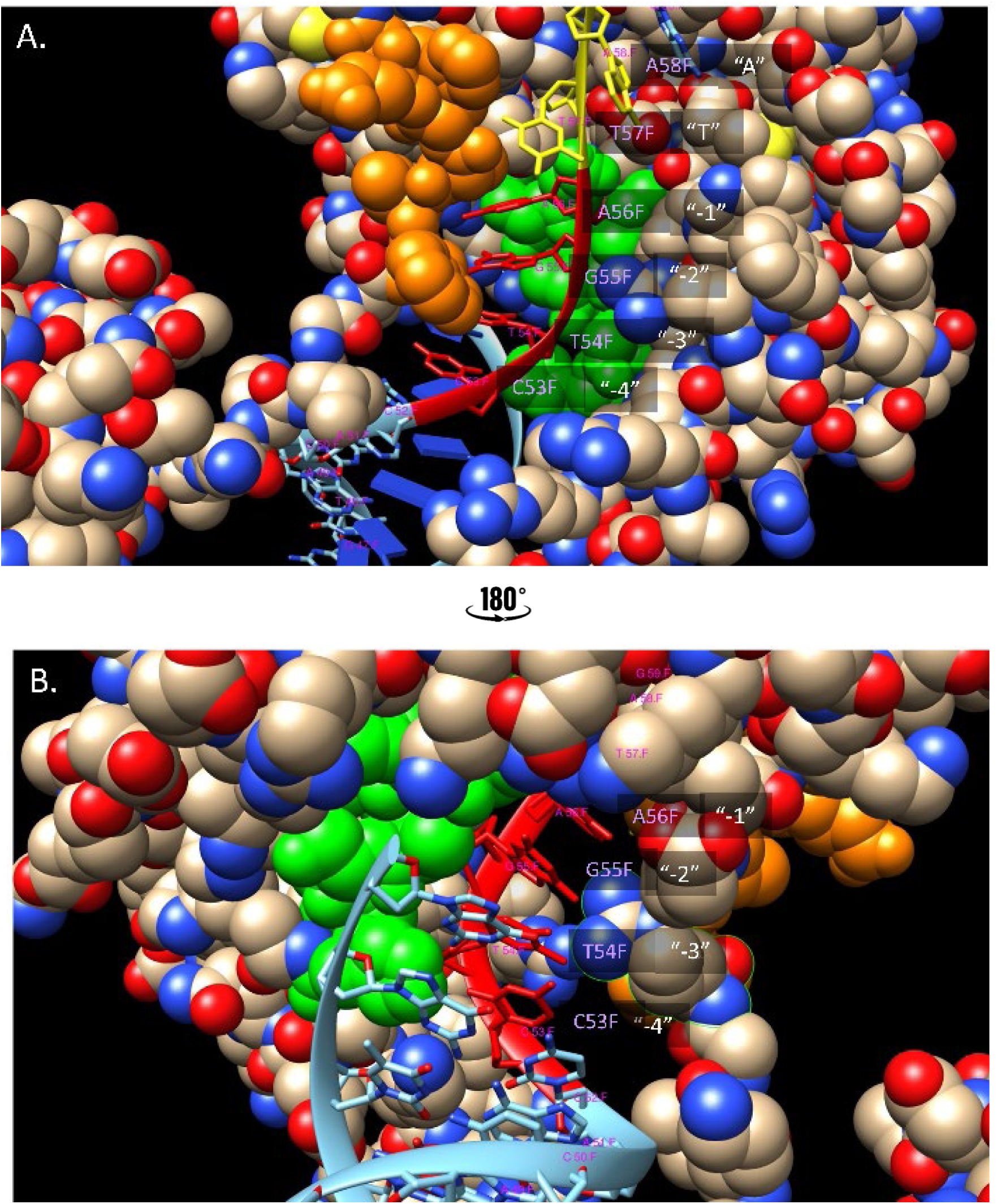
Crystal structure of complex between the Mos1 transposon and DNA. DNA double helix, with denatured (single strand) end in the pre-cleavage state. This is a stylized (cartoon) representation of the interaction. The red nucleotides represent C53-T54-G55-A56 (sites -4…- 1). The yellow nucleotides represent the TA site, T57-A58. The blue nucleotide is G59, which is site +1. The transposase itself is shown as a molecular surface. Amino acids 119-124 (WVPHEL) are colored orange and Asp284-His293 are colored green. The two images are vertical 180-degree rotations illustrating front and back views of the transposon-DNA interaction.

We demonstrated the utility of our model of nucleotide biases on Himar1 insertion frequencies by using it to improve gene essentiality predictions via the TTN-Fitness method. One way to determine the essentiality of a gene is to take the average count of insertions at all the TA sites in the gene, and determine the essentiality based on a set of cutoffs (Zomer, Burghout et al. 2012). This method treats all TA sites as being equivalent a priori (i.e., as independent, and identically distributed observations, with equal prior probability of insertion), and does not allow for site-specific differences that can greatly affect the insertion count at each site. Incorporating these surrounding nucleotides takes out (or corrects for) the effect of insertion biases and focuses the analysis on true biological effects, thus increasing our certainty in fitness calls for these genes. In the TTN-Fitness method, we fit a linear model to the insertion counts at TA sites, incorporating the gene in which it resides and the surrounding nucleotides each site as covariates. The coefficients associated with genes in the regression model reflect how much the mean insertion counts in the gene deviate from the global average, after correcting for the expected insertion counts at each site in the gene. For most genes, predicted fitness did not change substantially between the ablative Genes-only model and the Gene+TTN model of the TTN-Fitness method. However, the assessment for some notable genes did change with the inclusion of tetra-nucleotide features. PGRS13 was implied to be a ‘Growth Defect’ gene by the previous insertion count based methodology due to the low insertion counts at its TA sites. However, sites in the gene are surrounded by mostly ‘G’s and ‘C’s which have been determined by Himar1 preferences to suppress insertions. So, the insertions are low, but are expected to be low, and thus the gene is determined to be less essential than previously predicted. The Gene+TTN model used in the TTN-Fitness method has an advantage for small genes with less than or equal to 3 TA sites (220 in H37Rv genome) such as Rv3461c (*rpmJ*), previously undetermined by essentiality estimates. The model is less susceptible to noisy counts (high or low) at individual sites because we can compare the observed counts at those sites to expected counts from their nucleotide context, correcting for the effect of insertion biases, and thus improving the identification of conditionally essential genes and genetic interactions, i.e., to better distinguish true biological fitness effects by comparing the observed counts to expected counts using a site-specific model of insertion preferences. This method could also be helpful for analyzing differences in essentiality of genes between different strains (e.g. clinical isolates), where the TTN-Fitness model can correct for expected counts at TA sites to account for differences in the surrounding nucleotides (e.g. due to the different genetic backgrounds of the libraries).

## Methods and Materials

### Dataset of 14 independent Himar1 insertion libraries of *M. tuberculosis* H37Rv grown in vitro

We obtained 14 independent TnSeq libraries in *M. tuberculosis* H37Rv previously analyzed (DeJesus, Gerrick et al. 2017), representing a combined total of 35,314,576 independent insertion events by the Himar1 transposon. All libraries were treated uniformly, grown in standard laboratory medium (7H9/7H10). Every library in the 14 replicates has a mean saturation i.e., percentage of TA sites in the genome with 1 or more transposon insertions, of 0.65, totaling to a saturation of 0.85 for the entire dataset. As these are 14 independent libraries, the probability of a non-essential site with zero insertions for stochastic reasons is quite small. However, there is a lack of insertions in non-permissive sites in non-essential regions, which account for 9% of all TA sites. Most of the remaining sites with zero insertions correspond to essential regions.

This high level of saturation enabled us to reliably observe the nucleotide bias of insertion counts at different TA Sites. The dataset was normalized using TTR normalization in Transit (dividing by the total counts in each dataset, with top 1% trimmed to mitigate influence of outliers, and scaling back up so the mean count at non-zero sites is 100.0). We identified essential regions as consecutive sequences of 6 or more TA sites with counts of two or less and subsequently removed them. Using the resulting dataset, we were able to explain nucleotide bias at TA Sites not only for H37Rv but also for other mycobacteria and non-mycobacterial Himar1 TnSeq datasets.

### Significance of the Correlation of Insertion Counts between TnSeq Datasets

The correlation of insertion counts at TA sites between TnSeq datasets was calculated using a Pearson correlation coefficient. As mentioned previously, the log of insertion counts was used, since the Pearson Correlation Coefficient assumes that the input data is normally distributed. The two-tailed T-test for the means of two independent samples was used to measure whether the expected value differs significantly between samples. Since we do not assume population variance between the two datasets is equal, the Welch’s t-test is performed.

### 10-Fold Cross Validation Linear Regression

The data was split for 10-Fold cross validation using sklearn.model_selection.KFold. Within these folds, we used sklearn.Ridge with alpha=0.1 to train and test linear models (target values of log insertion counts or LFCs) with L2 regularization.

### Hyper parameter Tuning the Neural Network

The data was separated into training and testing using a 70-30 train-test split. We used 10-Fold cross validation on the 70% training split of the data to tune the number of nodes per hidden layer, number of hidden layers, the activation function, value of alpha and whether to use early stopping. We used scikit-learn’s GridSearchCV to perform this operation and checked the accuracy of the final hyper parameters set on the reserved 30% set. Afterwards, we used the tuned parameters to perform a 10-fold cross validation on the whole dataset to accurately judge the model and account for data biases.

### Mean LFCs per Nucleotide-Position pair

For every position ±20 bp from the TA site, we filtered for nucleotides ‘A’, ’T’, ’C’, and ’G’. We took the mean LFC of the training samples (TA sites) with that nucleotide in that position. This calculation yielded the mean LFC for each nucleotide at each position 20 bp from the TA site, which were then visualized as a heatmap with a diverging color palette.

### Model Adjustment Calculations

Each TnSeq dataset has a slightly different LFC distribution. Thus, the predictions of a TnSeq dataset from the STLM, trained on H37Rv data, had to be adjusted. This was accomplished by a simple regression-based procedure. First, we determined the linear relationship between the mean LFC for each tetra-nucleotide in H37Rv by regressing it against the mean LFC of each tetra-nucleotide in our target strain. The linear relationship could be represented as *targetStrainLFCs* = *m* * *H*37*RvLFCs* + *offset*. We used this relationship to adjust the LFC predictions made by the STLM using the target strain’s *LFC_adjusted_* = *m* * *LFC_STLM_* + *offset*.

### Average Change in Observed LFC vs. Average Change in Predicted LFC Between Strains

In comparing the genome sequences of *M. abscessus* ATCC 19977 and the Taiwan49 clinical isolate, there are 9,303 TA sites with exactly one SNP in the surrounding ±4 bp window. There are 8 positions and 12 possible substitutions per position, thus 96 possible SNPs that can occur. For each of these possible nucleotide changes, we calculated the difference between the observed LFC in the reference strain and the observed LFC in the isolate strain. The mean of this difference was determined to be the mean observed LFC difference for that SNP. We performed a similar calculation for the predicted LFCs. Using the STLM, we found the predicted LFCs at a TA sites in the reference strain and predicted LFCs at TA sites with a specific SNP in the clinical isolate. The average of the difference in these two predicted LFCs was the mean change in predicted LFC.

### Using the Bernoulli Distribution to filter small essential genes before training the Gene+TTN model

The first step in fitness estimation is to identify and remove any essential genes. These genes are excluded from the Gene+TTN analysis. First, larger essential genes (with > ∼10 TA sites) are identified using the Gumbel method in TRANSIT. Then smaller essential genes with no insertions are identified and removed based on a Bernoulli calculation. Given the probability that an insertion does not occur (*p*=1.0- saturation), the probability of *k* TA sites out of *n* total having no insertions follows the Binomial Distribution:

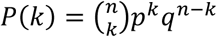

Thus, the probability that all TA sites in a gene have 0 insertions is a Bernoulli distribution where *k=n*:

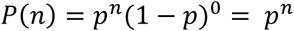

We use this formula to determine the minimum *n* such that *P(n)*<0.05, and we then label any genes with *n* or more TA sites, all of which have insertion counts of 0, as ‘Essential-B’ (“ESB”). This method is a necessary additional step to the Gumbel method to find smaller genes that may have been missed, especially in datasets with lower saturation.

### Availability

The source code (Python scripts) for performing the calculations described in this paper (including the TTN-Fitness model) are available at github.com/ioerger/TTN-Fitness. The raw data files (wig files with insertion counts at TA sites) for the 14 replicate libraries of *M. tuberculosis*, along with 3 replicates for *M. abscessus* Taiwan49, can also be found in the demodata/ directory of the same github repository.

## FIGURES and TABLES

**Supplemental Figure S1:**
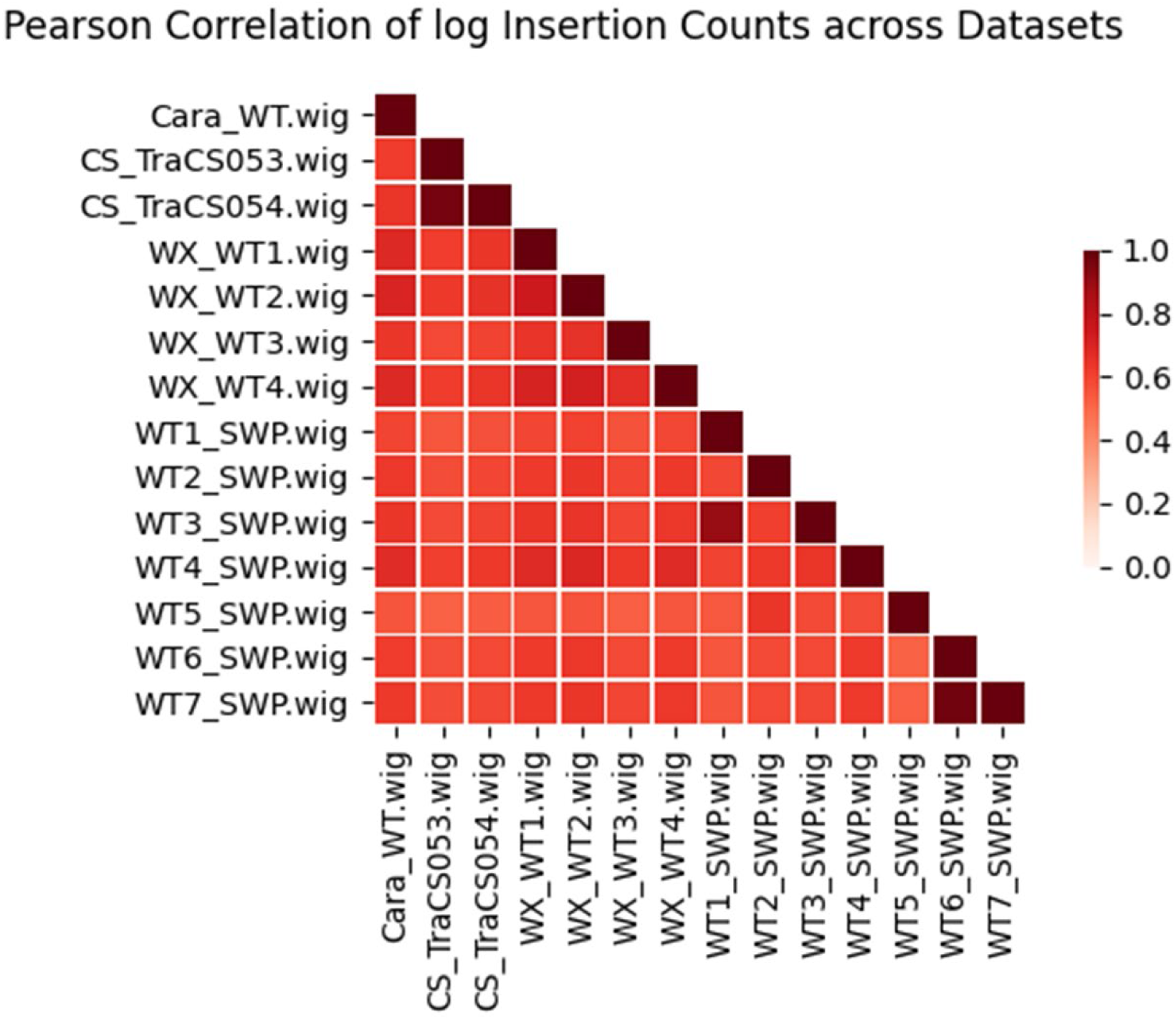
Heatmap of Correlation of Wig Files. The correlation of log insertion counts in the 14-replicates wig files using the Pearson correlation coefficient as recorded in Supplemental Table T1.

**Supplemental Figure S2:**
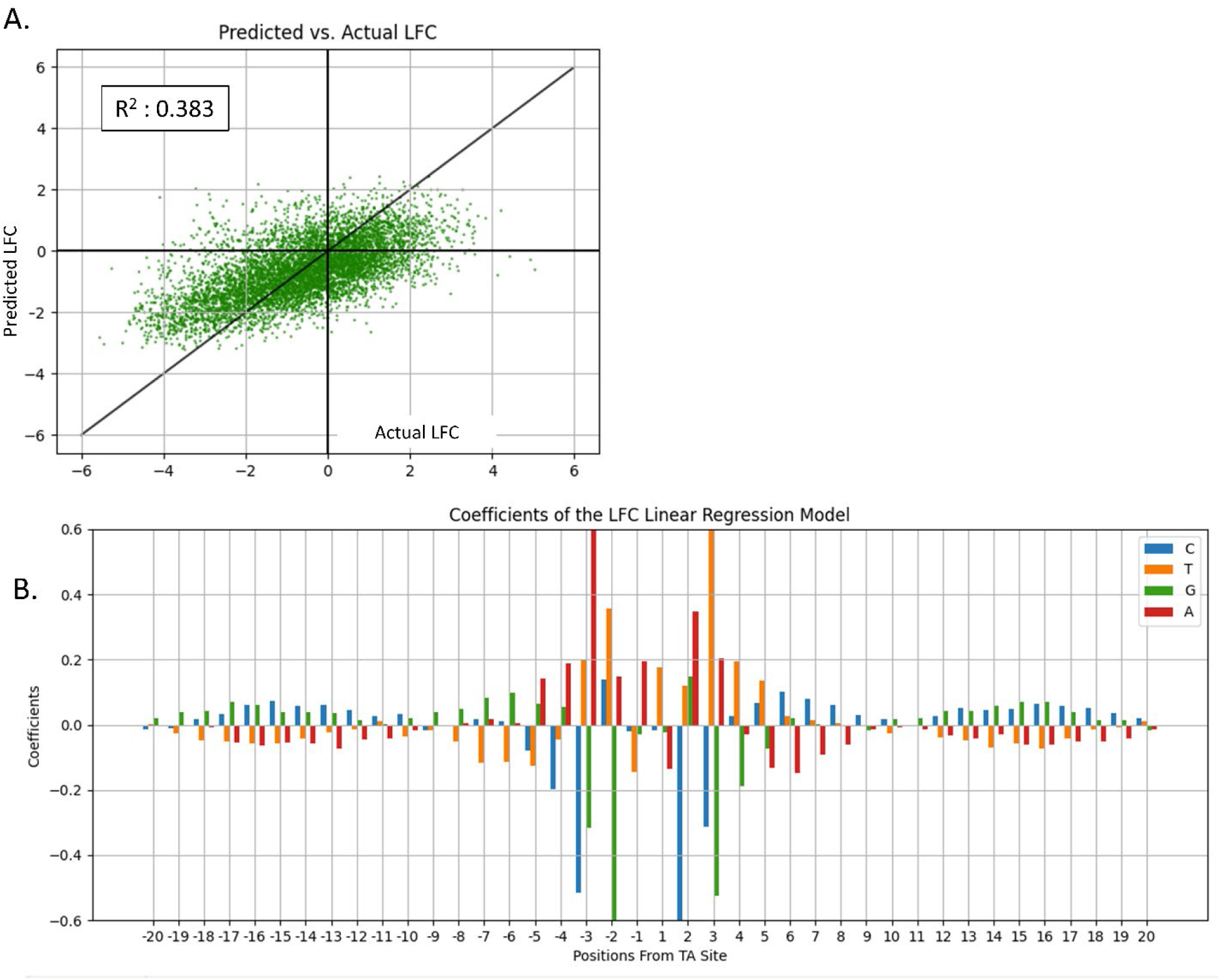
Coefficients from Linear Model Trained using nucleotides in all 40 positions to predict LFCs. Panel A. shows Predicted Counts vs. Actual LFC using Linear Regression. The average predictive power of the linear regression model trained with one-hot-encoded nucleotides in 20 bp from the TA site as the input and LFCs as the output using 10-fold cross validation. The predictive power was not much higher than the previous Insertion Counts model, but the variance has decreased, indicating a better model. **Panel B** shows coefficients from the trained model. The coordinates along the x-axis give the positions relative to, but not including, the TA site. A symmetric pattern is visible in positions -4,-3,-2,-1 and +1, +2, +3, +4. The non-permissiveness pattern (CG)GnTAnC(CG) is visible in this window as well as high coefficients associated with “A” and “T”.

**Supplemental Figure S3:**
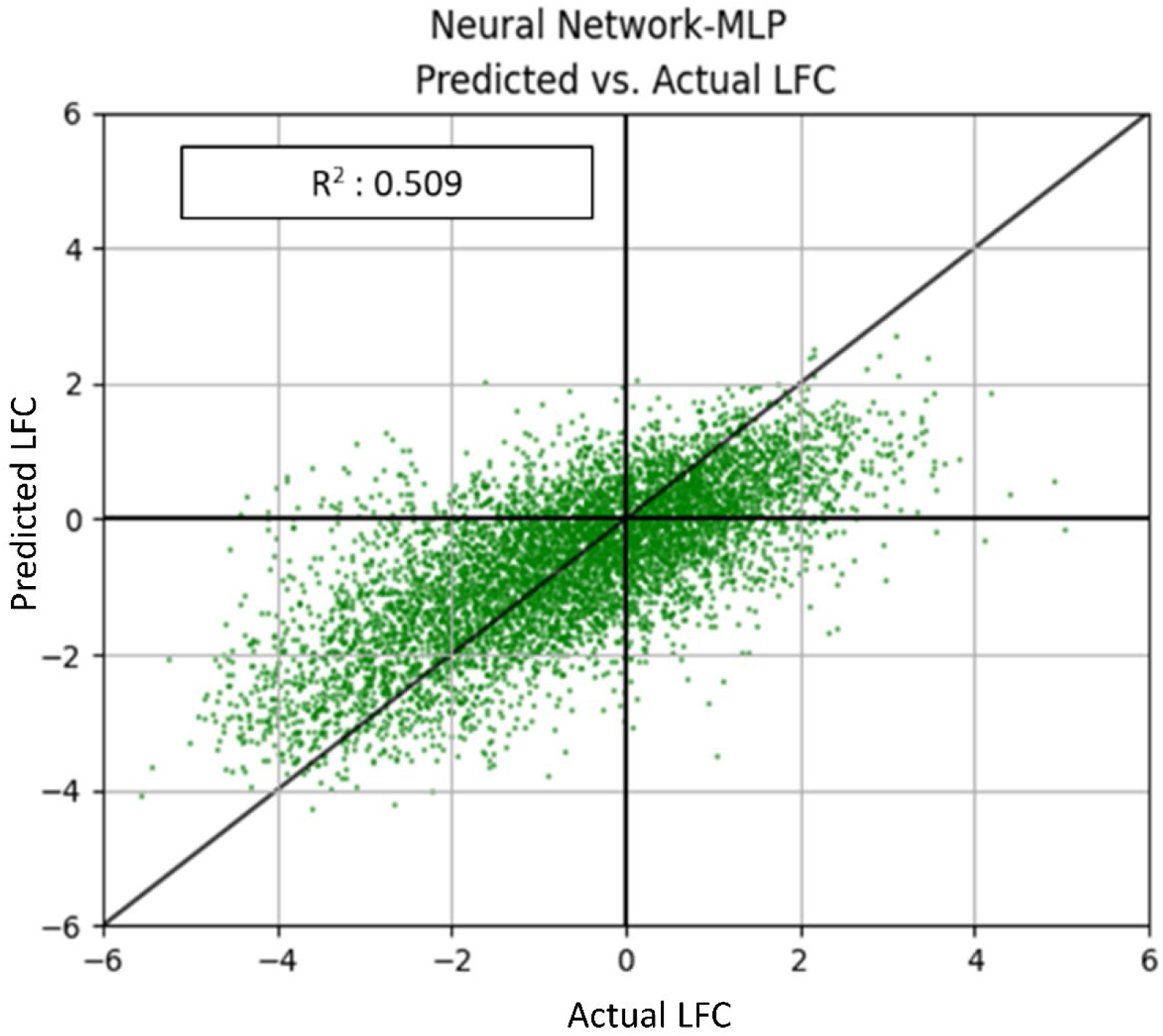
Predicted LFC vs. Actual LFC using Feed Forward Neural Network. The input to this linear model was all the one-hot-encoded nucleotides, and the target value as the LFCs. Using 10-fold cross validation on 70% of the dataset, we found the ideal parameters: ’activation’: ’tanh’, ’alpha’: 0.05, ’early_stopping’: True, ’hidden_layer_sizes’: (100,), ’learning_rate’: ’constant’, ’max_iter’: 500, and ’solver’: ’adam‘. We tested these hyper-parameters on the remaining 30% of the test data and got a fairly high performing model. We applied these hyper parameters and assessed the model’s fit to the data by performing a 10-fold cross validation of the entire dataset. This yielded an average predictive power (i.e. R^2^) that was higher than the previous Insertion Counts Model and LFC Model and the variance has decreased, indicating a better model.

**Supplemental Figure S4:**
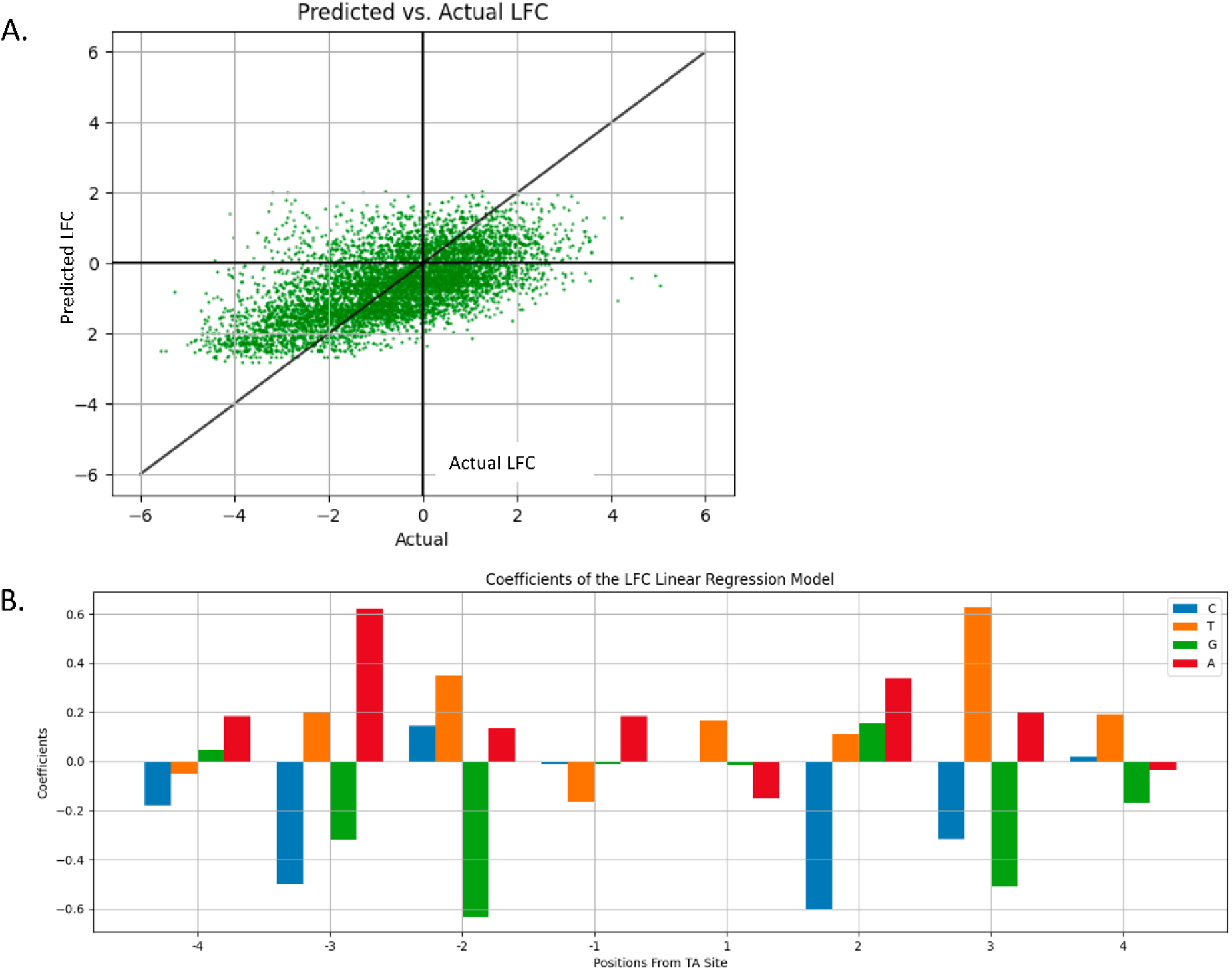
LFC Prediction using linear regression with only nucleotides in -4…+4 positions from the TA site. **Panel A** shows Predicted Counts vs. Actual LFC using Linear Regression. The average predictive power of the Linear Regression Model trained with one-hot-encoded nucleotides in 4 bp from the TA site as the input and LFCs as the output using 10-fold cross validation. The predictive power is moderate (R^2^=0.352), meaning it can explain 35% of the variation in insertion counts based on surrounding nucleotides, not much different than the LFC linear model trained using all 40 nucleotides, indicating the nucleotides in this window are very important. **Panel B** shows Coefficients from the trained model. The coordinates along the x-axis give the positions relative to, but not including the TA site. These coefficients are almost identical to the relative magnitudes of the nucleotides in the -4…+4 window of LFC linear model and the log insertion count linear model.

**Supplemental Figure S5:**
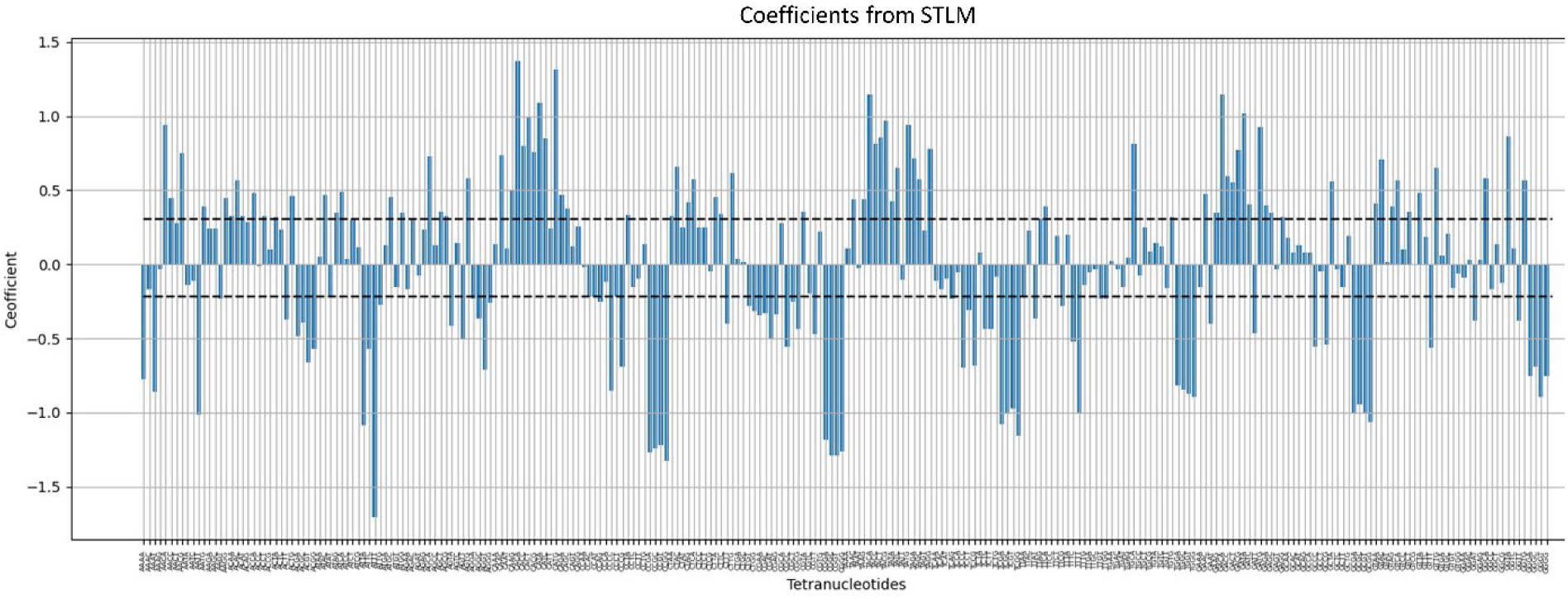
Coefficients from the STLM. Coefficients of a linear model trained on one-hot-encoded TTN and a target value of LFCs. The highest values are for tetra-nucleotides XAXX and the lowest are XCXX and XGXX, where X is any nucleotide A, C, T or G. P-values obtained from the Wald test, with FDR adjustment showed that coefficients ≤ -0.234 and ≥ 0.315 are significant.

**Supplemental Figure S6:**
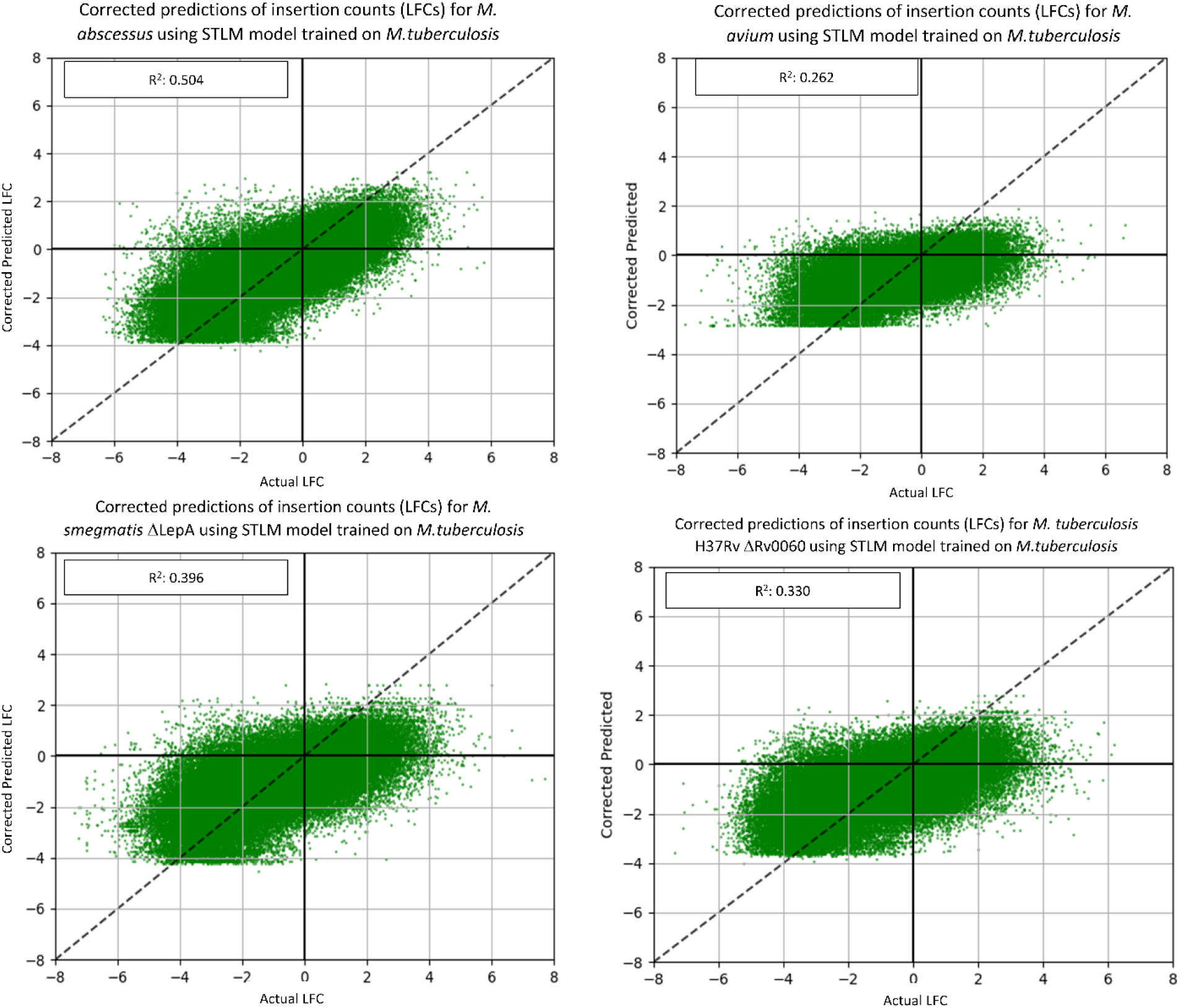
Predictive Power of STLM on Mycobacterial Datasets. The predictive power of the STLM on the Mycobacterial datasets have been varied. However, a R^2^ value greater than 0.25 for nearly all the datasets indicates that the nucleotide biases explain *at least* a fourth of the variance in insertion counts with nucleotide biases.

**Supplemental Figure S7:**
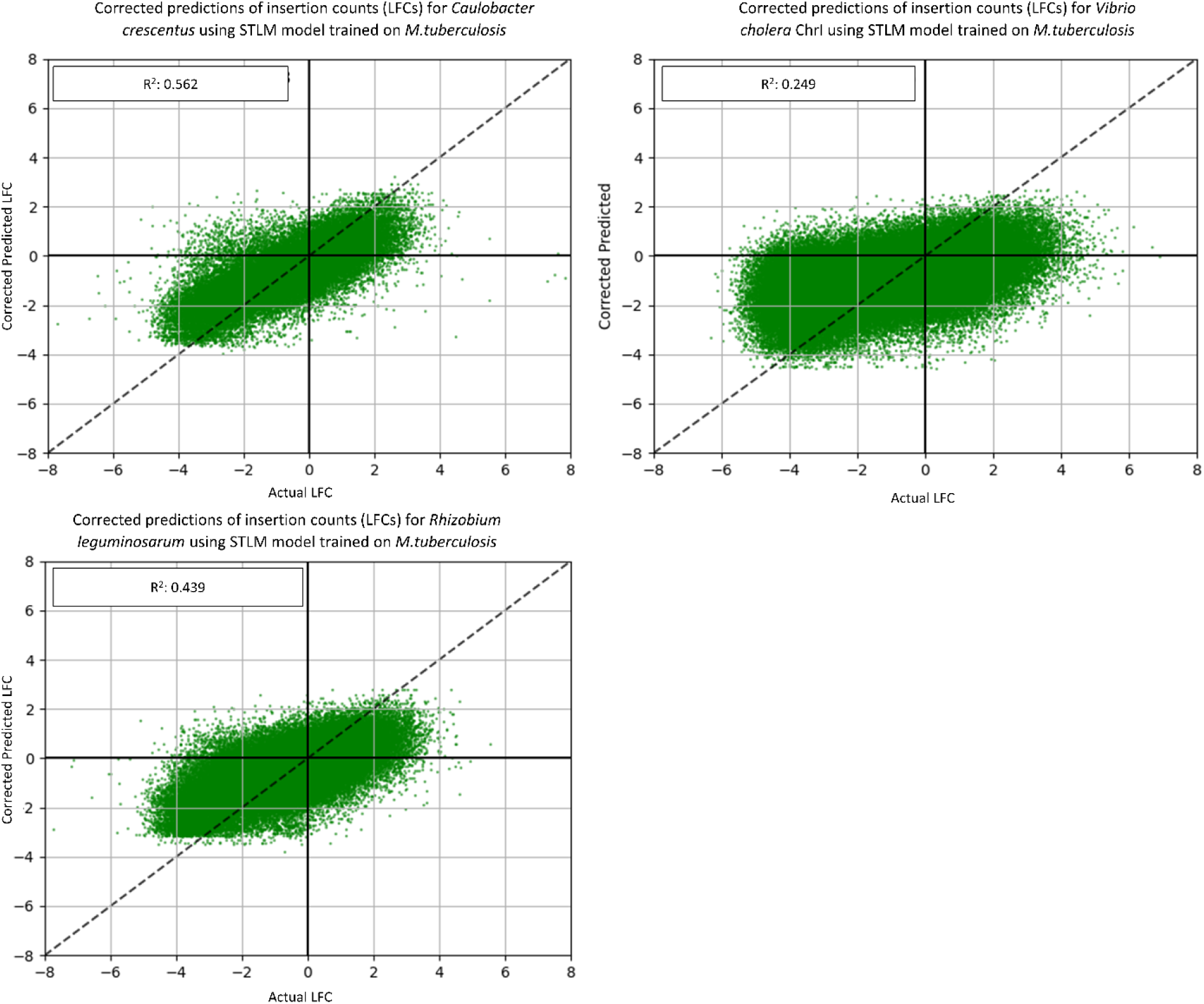
Predictive Power of STLM on non-Mycobacterial Datasets. The predictive power of the STLM on non-Mycobacterial datasets have been more varied than the Mycobacterial datasets. *Caulobacter* has a high R^2^ value, whereas *Vibrio* has quite a low R^2^ value. However, a R^2^ value greater than 0.10 for nearly all the datasets indicates that the nucleotide biases explain at least *some* of the variance in insertion counts with nucleotide biases.

